# Meta-atlas of Juvenile and Adult Enteric Neuron scRNA-seq for Dataset Comparisons and Consensus on Transcriptomic Definitions of Enteric Neuron Subtypes

**DOI:** 10.1101/2024.10.31.621315

**Authors:** Joseph T. Benthal, Aaron A. May-Zhang, E. Michelle Southard-Smith

## Abstract

**Background:** The enteric nervous system (ENS) is a complex network of interconnected ganglia within the gastrointestinal (GI) tract. Among its diverse functions, the ENS detects bowel luminal contents and coordinates the passing of stool. ENS defects predispose to GI motility disorders. Previously, distinct enteric neuron types were cataloged by dye-filling techniques, immunohistochemistry, retrograde labeling, and electrophysiology. Recent technical advances in single cell RNA-sequencing (scRNA-seq) have enabled transcriptional profiling of hundreds to millions of individual cells from the intestine. These data allow cell types to be resolved and compared to using their transcriptional profiles (“clusters”) rather than relying on antibody labeling. As a result, greater diversity of enteric neuron types has been appreciated. Because each scRNA-seq study has relied on different methods for cell isolation and library generation, numbers of neuron clusters and cell types detected differs between analyses. Cell counts in each dataset are particularly important for characterization of rare cell types since small numbers of profiled cells may not sample rare cell types. Importantly, each dataset, depending on the isolation methods, may contain different proportions of cells that are not detected in other datasets. Aggregation of datasets can effectively increase the total number of cells being analyzed and can be helpful for confirming the presence of low-abundance neuron types that might be absent or observed infrequently in any single dataset.

**Results:** Here we briefly systematically review each *Mus musculus* single cell or single nucleus RNA-sequencing enteric nervous system dataset. We then reprocess and computationally integrate these select independent scRNA-seq enteric neuron datasets with the aim to identify new cell types, shared marker genes across juvenile to adult ages, dataset differences, and achieve some consensus on transcriptomic definitions of enteric neuronal subtypes.

**Conclusions:** Data aggregation generates a consensus view of enteric neuron types and improves resolution of rare neuron classes. This meta-atlas offers a deeper understanding of enteric neuron diversity and may prove useful to investigators aiming to define alterations among enteric neurons in disease states. Future studies face the challenge of connecting these deep transcriptional profiles for enteric neurons with historical classification systems.

## BACKGROUND

The enteric nervous system (ENS) is a network of ganglia within two layers of muscle within the gut wall that allows for passing of stool via peristalsis. The ENS, like the central nervous system and brain, is made up of several distinct neuronal subtypes whose function can correspond to their positioning along the intestine.^1^ Historically, the ENS and its cell types have been defined by several techniques, including dye-filling, immunohistochemistry, retrograde labeling, electrophysiology, and numerous other methods.^2^ From these studies, the ENS field has gained a broader understanding of how the cells of the ENS function *in situ* and *in vitro*, allowing researchers to model perturbations in ENS function, which often result in gastrointestinal (GI) motility disorders in patients. It has been shown via some of these methods that the balance of subtypes of enteric neurons are altered in some GI motility disorders, including Hirschsprung disease.^3,4^ Given the imbalance of enteric neuronal subtypes in some GI motility disorders, it is important for treatment of these disorders that we understand the diversity of cell types in the ENS so that patients might have effective treatments.

Advances in single cell transcriptomics have enabled profiling of individual cells from an array of tissues including the ENS where efforts to define enteric neuron diversity previously relied on immunohistochemistry, dye-filling, or pharmacological studies of single neurons. To date, at least 13 distinct mouse ENS single cell or nucleus datasets have been produced from 11 different manuscripts, with each publication using different isolation methods, strains/ages of mice, and methods of cell or nuclei isolation.^5–17^ Multiple reviews have been published extensively comparing these datasets.^2,18,19^ Each study profiling ENS progenitors and mature enteric neurons applied distinct approaches both technological and bioinformatic to produce single cell or nucleus RNA-seq data. Variables included differing mouse lines, ages, tissue dissociation methods, techniques for encapsulation/library production, and depth of sequencing.^5–17^ These differences have led to variation in the number of cells sampled, cell types detected, and genes whose expression labels distinct cell clusters.^2,18,19^ A main point of discussion in the ENS field centers on reaching consensus regarding how many ENS cell types exist and what methods should be prioritized for classifying and naming them for consistency across the literature. Combining enteric neuron scRNA-seq datasets in a strategic manner is one means to gain greater understanding of diversity of enteric neurons. In this work, we first systematically review each published single cell or single nucleus RNA-seq dataset derived from the ENS, highlighting the main contributions to the field that each work produced. We then make the case below for combining the more mature enteric neuron datasets and illustrate the integration process, outlining both advantages and caveats for this method in the attempt to transcriptionally define enteric neuron states. Our goal is to demonstrate how meta-analysis (the process of pooling independent datasets together into one larger aggregate) can yield consistent cell states across datasets, a higher sample size of enteric neurons and therefore greater resolution to detect rare cell states. We offer this resource of similarly processed datasets for future mining by the ENS community.

## METHODS

We briefly review ENS publications that utilize single cell or single nucleus RNA-sequencing. We then highlight how integration of single cell RNA-seq datasets can contribute to the field. We subsequently extract the relevant publicly available mouse enteric neuron single cell or nucleus RNA-seq datasets, reprocess, integrate, and perform analyses to deepen transcriptional definitions of enteric neuron subtypes.

### Literature Review

We collected all relevant publications of which we were aware and used those as a starting point for review. We also utilized Google Scholar and the Vanderbilt University Library website interface to search for and access other papers using various related search terms, including “ENS”, “Enteric Nervous System”, “scRNA-seq”, “single cell”, and “single cell RNA-sequencing”. We then screened papers for scRNA-seq datasets and excluded papers that were not peer-reviewed (e.g. preprint servers).

### Data Access and Downloading ENS scRNA-seq Datasets

We downloaded the data from the following sources. Wright and colleagues’ 47 to 52 days old mouse distal colon myenteric plexus snRNA-seq data was downloaded from Gene Expression Omnibus accession number GSE156905 in the form of matrix, barcode, and feature (gene) files.^11^ Morarach and colleagues’ P21 juvenile scRNA-seq mouse data were downloaded from Gene Expression Omnibus accession number GSE149524 in the form of matrix, barcode, and feature (gene) files.^10^ May-Zhang and colleagues’ 6-week-old mouse enteric neuron snRNA-seq data was downloaded from Gene Expression Omnibus accession number GSE153202 in the form of matrix, barcode, and feature (gene) files.^9^ We downloaded Zeisel and colleagues’ processed ENS P19, P20, and P21 scRNA-seq data in the form of a .Loom file from mousebrain.org.^12,20^ Drokhlyansky and colleagues’ 11 to 104-weeks-old mouse enteric neuron data were downloaded (each cell type downloaded separately) from the Single Cell Portal at the Broad Institute website at accession number SCP1038), which requires a Google login and the data were in the form of matrix, barcode, and feature (gene) files.^5^

### Dataset Reprocessing

Each dataset’s individual sc/snRNA-seq library was read into R using Seurat’s Read10x and CreateSeuratObject functions with a minimum of 3 cells and minimum of 200 features.^21–25^ We then used Seurat’s PercentageFeatureSet function to get the percentage of mitochondrial gene expression per cell based on the gene prefix pattern “∧mt-“. We then plotted histograms, violin plots, and FeatureScatter (Seurat plotting function) plots of percent mitochondrial gene expression, nFeature_RNA, and nCount_RNA. These were used in choosing quality control (QC) metrics when filtering the data. We also considered the QC metrics and clustering parameters listed in the publications. Please see the code for each dataset reprocessing for the specific metrics, which ranged from 200-2000 genes per cell/nuclei as the lower bound (nFeature_RNA > 200-2000). We decided that 20 percent mitochondrial RNA per cell in each dataset was of sufficient quality (percent.mt < 20). We note that the RAISIN-seq dataset from Drokhlyansky et al. have extremely high values of nCount_RNA at a mean of just under 1,000,000 (2020). This may have to do with the method (SMART-Seq2) that was used for assaying these “RAISINs”.^5,26^ After QC filtering, we applied Seurat’s SCTransform v2 to each run, regressing out percent mitochondrial genes.^27,28^ Next, we ran principal component analysis (PCA), uniform manifold approximation projection (UMAP) dimensionality reduction, Seurat’s FindNeighbors, and FindClusters. After finding clusters and visualizing the UMAP through Seurat’s DimPlot function, we used DoubletFinder to identify and filter putative heterotypic doublets.^29^ This requires an expected percentage of doublets, which can be estimated based on the number of cells loaded into a 10X Chip.^30^ We used the number of cells in the object to estimate the cells recovered along with a linear function based on data found on the 10X Genomics website to estimate the multiplet rate.^31^ This rate was used in the round function of DoubletFinder. SCTransform v2, RunPCA, RunUMAP, FindNeighbors, and FindClusters were then run again on the data filtered for putative heterotypic doublets. After running each run of the datasets through this pipeline, we then integrated the runs using SCTransform v2.^28^ For most datasets, we reduced the number of clusters for Figure 2A-F by reducing the clustering resolution parameter. See the GitHub code and our Open Science Framework page for more details, where we outline code for the estimated doublet rate linear function and provide RMD files for each dataset. Code for reducing the cluster number for most datasets is in a separate R code file.

### Integration of datasets and Meta-Atlas Analysis

First, metadata were added to each dataset object generated, including the dataset first author last name, year of publication, age mouse, tissue section, etc. After, datasets were integrated and batch corrected via Seurat V5 by sequencing run into a meta-atlas using SCTransform v2 per the Seurat tutorial (https://satijalab.org/seurat/articles/seurat5_integration).^21–25,27,28^ We integrated by run to control for both the tissue segment and the age of the mice at the time of neuron isolation to the best of our ability. After integration and batch correction via SCTransform v2, we processed the meta-atlas dataset with RunPCA, RunUMAP, FindNeighbors, and FindClusters. Next, we normalized and scaled the “RNA” assay using Seurat’s NormalizeData and ScaleData functions. We used the PrepSCTFindMarkers function to prepare the data for differential gene expression for each cluster (using Seurat’s FindAllMarkers function). We then ran FindAllMarkers with both the SCT-corrected data and the normalized RNA data, for which the differences were minimal. We used the RNA-assay-based FindAllMarkers results going forward. We then displayed the top 30 putative marker genes per cluster (sorted by Log2FoldChange and significant Bonferroni-adjusted p-value from FindAllMarkers Wilcoxon rank-sum test) in a heatmap to see clustering patterns via gene expression. To reduce the occurrence of multiple clusters sharing similar expression patterns, we reran the clustering algorithm with a lower resolution. After re-running FindAllMarkers and re-generating the heatmap, the top 30 enriched genes for each cluster showed much less overlap than was originally observed. We then visually examined plots generated from the top 30 FindAllMarkers results for each cluster and chose genes that were unique to each cluster.

Because of the lack of specific expression of putative marker genes in the meta-atlas clusters for the subset juvenile datasets, we opted to reintegrate the data using the same approach outlined above including only the adult datasets. The adult meta-atlas only included runs from May-Zhang et al., 2021, Drokhlyansky et al., 2020, and Wright et al., 2021.^5,9,11^

Morarach et al., 2021 found that there was independent expression of *Etv1* and *Bnc2* in each of their developing “branched” neuronal populations that is maintained into their juvenile stages, yet some clusters at in their juvenile scRNA-seq data do not express either of these markers.^10^ To perform differential gene expression on *Etv1* and *Bnc2* expressing clusters versus clusters that do not express these, as well as finding genes that mark new subclusters, we performed differential gene expression via Seurat’s FindMarkers (Wilcoxon rank-sum test) function for these two groups. The top genes (sorted by Log_2_FoldChange and significant Bonferroni-adjusted p-value) were then assessed for specificity. We used the same approach when performing differential gene expression between subtypes of enteric neurons based on unsupervised clustering. The marker genes identified in the figures were found by visual inspection for exclusivity for a specific cluster across all datasets via Seurat’s FeaturePlots. We used the same approach when performing FindMarkers between cell subtypes, which is outlined further below. These were used for Figures 5C’ and 5E’.

To perform differential gene expression analysis by sex, we subset the adult data further to only include sex annotations (subtracting Wright et al., 2021). We performed this analysis both by cluster using the Wilcoxon rank-sum test and via Seurat’s Logistic Regression method by cluster per segment, regressing out the dataset label metadata (i.e., first author plus the year of publication) to account for dataset differences.

### Availability of Data and Materials

The datasets and R code supporting the conclusions of this article are available in the Open Science Framework repository, https://osf.io/evjx4/?view_only=38c1fb85b84e486db9f8c26ea5c61e65.

## RESULTS

### A multitude of neuron types revealed by single cell RNA-sequencing studies

Advances in single cell transcriptomics have enabled profiling of individual cells from an array of tissues including the ENS, where efforts to define enteric neuron diversity previously relied on immunohistochemistry, dye-filling, or pharmacological studies of single neurons. To date, at least 13 distinct mouse ENS single cell or nucleus datasets have been produced from 11 different manuscripts, with each publication using different isolation methods, strains/ages of mice, and methods of cell or nuclei isolation.^5–17^ Multiple reviews have been published extensively comparing these datasets.^2,18,19^ Here, we first review the dataset features and major findings of recent single cell/nuclei profiling studies as a prelude to our main goal of describing how data aggregation can be undertaken using the ENS as an example.

Lasrado and colleagues produced the first ENS scRNA-seq dataset.^7^ The study aimed to define spatial coordination of murine ENS development by relying on a *Sox10-creER^T^*^2^ transgenic activating a ROSA confetti reporter at 12.5 days post coitus (dpc). The analysis identified spatially distinct bipotent and fate-restricted progenitors/precursors originating from neuronal cells with limited proliferative capacity and glial progenitors with higher proliferative capacity. Using fluorescence activated cell sorting (FACS) the team isolated confetti-tagged ENS cells at 12.5-13.0 dpc for scRNA-seq libraries. While these experiments produced a small data set containing 120 cells, the analysis identified three cell clusters distinguished by expression of typical glial (*Erbb3*, *Sox10*, *Fabp7*), progenitor (*Plp1)* and neuronal (*Tubb3*, *Elavl4*, *Ret*) marker genes. The expression profiles were consistent and further refined earlier reports of neuronal and glial lineage divergence during colonization of the fetal mouse intestine and offered an initial view of the genes being transcribed at this critical stage in ENS development.^32^

A second ENS scRNA-seq dataset was part of a larger mouse nervous system atlas from Zeisel et al., 2018, in which mouse ENS cells were collected from postnatal (P) stages.^12^ This study relied on lineage tagging of ENS populations from *Wnt1-Cre;ROSA26tdTomato* reporter mice for fluorescent postnatal labeling of neural crest derivatives at P19, P20, and P21 (Table 1). Generation of tissue dissociates from younger mice is easier than in older animals, and the ages collected were designed to profile changes in the ENS that occur around the time of weaning (P20-P21) when animals transition to solid food. In this effort, analysis of the resulting scRNA-seq libraries found most transcriptional profiles originated from enteric glia. This outcome most likely resulted from loss of membrane integrity among neurons during tissue dissociation that shears neuronal processes and allows uptake of viability dyes leading to cells being rejected during flow sort isolation. Despite this challenge, the team successfully profiled 727 enteric neurons that resolved into nine cell clusters in t-SNE representations using typical clustering algorithms. Overall, the analysis found gene expression patterns typical of glutamatergic, cholinergic, and nitrergic enteric neuron types.

**Table 1.**
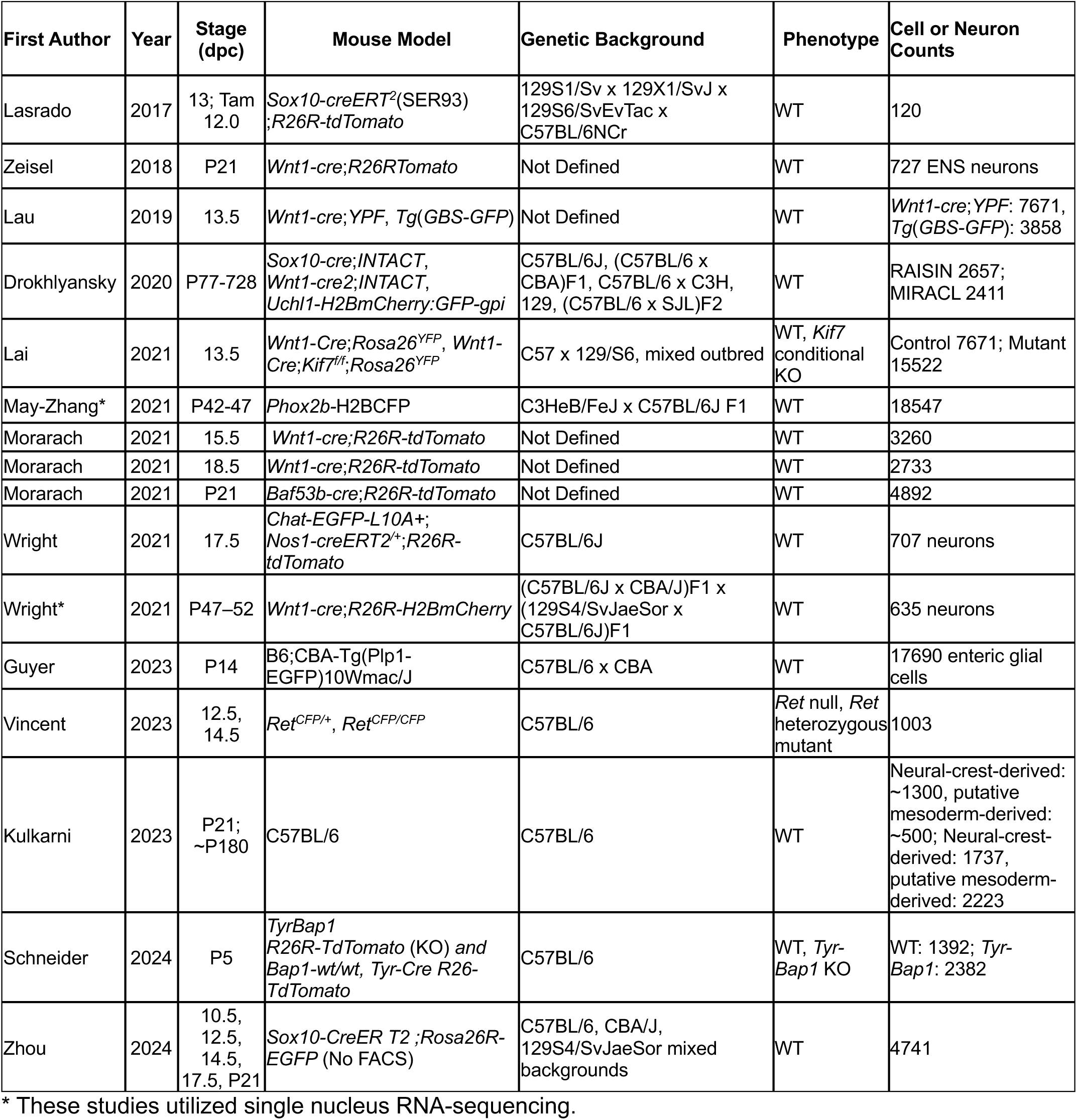
Characteristics of ENS RNA-sequencing datasets.

Lau and colleagues subsequently produced an extensive fetal mouse ENS scRNA-seq data set at 13.5 dpc while analyzing Hedgehog signaling.^6,8^ This team also relied on the *Wnt1-cre*;Rosa26*YFP* to isolate 7671 neural crest cells from fetal mice. They compared their mouse transcriptional profiles to scRNA-seq from human pluripotent stem cell (hPSC)-derived neural crest derivatives. From principal component analysis, the group identified 8 mouse and 12 human cell clusters. Like the Lasrado study, Lau and colleagues also observed two distinct differentiation paths for enteric neurons and glia. They then captured additional enteric lineages for scRNA-seq by utilizing a GLI1-3 GFP fluorescent reporter mouse line, *Tg*(*GBS-GFP*), that mirrors hedgehog signaling levels. From this effort, they produced transcriptional profiles from 2017 Hedgehog “on” and 1841 Hedgehog “low” signaling cells. Integration of these cells into their original 7671-cell scRNA-seq dataset found these cells are distributed across all 8 mouse cell clusters, which suggests GLI1-3 activity is dynamic across ENS differentiation.

Optimization of cell isolation methods and new informatic tools led to two independent studies in quick succession that compared scRNA-seq data from adult ENS of mice and humans authored by Drokhlyansky et al and May-Zhang et al.^5,9^ Drokhlyansky and colleagues transcriptionally profiled adult mouse and human ENS from older ages at single cell resolution.^5^ This team relied on two complementary strategies that facilitated transcriptional profiling of enteric neurons via single nucleus RNA-Seq (snRNA-Seq). The first strategy consisted of a new technique for isolation of nuclei with bound ribosomes on the outer nuclear membrane called Ribosomes and Intact single nucleus (RASIN) RNA-Seq that generated sequence data with higher spliced mRNA recovery than RNA-Seq of nuclei alone. The second strategy, Mining Rare Cells sequencing (MIRACL-seq), informatically selected droplets containing rare cells from overloaded single cell encapsulations generated from human bowel. For comparison of human enteric neurons with those of mice, the team isolated ENS cells from adult mice of both sexes ranging from 11 to 104 weeks of age. Transgenic lines that fluorescently tagged nuclei of neural crest lineages (*Sox10-cre:INTACT, Wnt1-cre:INTACT*) or enteric neurons (*Uchl1-H2BmCherry:GFP-gpi*) were used for flow sorting to enable transcriptional profiling of 2657 neurons and 3039 glia from colon. The authors identified colonic neuron clusters that putatively represent 21 distinct neuron types consisting of excitatory and inhibitory motor neurons, interneurons, sensory types, and secretomotor/vasodilator neurons as well as three enteric glial types. The analysis indicated regional differences in the distribution of enteric neuron types in the mouse colon that also varied by age and circadian phase. The Drokhlyansky team then applied MIRACL-seq to characterize mouse and human enteric neurons. This strategy captured transcriptional profiles of 1938 mouse neurons that parsed into 18 neuron types with some notable differences detected between colonic and ileal neurons. Similar MIRACL-seq profiling of muscularis propria from human colon cancer patients ranging in age from 35-90 years identified 1,445 neurons that segregated into 14 neuron classes. By comparing the mouse RNA-seq data to the human MIRACL-seq profiles the authors concluded that multiple types of enteric neurons are transcriptionally similar between species.

Comparison of transcriptional profiles between species and subsequent *in situ* localization of neuronal subtype markers in mice and humans performed by May-Zhang and colleagues reached slightly different conclusions.^9^ This team aimed to compare enteric neuron gene expression between healthy young adult humans (18-35 years of age) and mice (6-7 weeks of age). Their efforts initially focused on capture of myenteric neuronal nuclei from duodenum, ileum, and colon based on *Phox2b*-H2BCerulean transgene expression that is highly expressed in differentiated neurons.^33^ The team successfully captured more than 25,000 nuclei with retention of 18,500 profiles after quality control criteria were applied. Clustering of these data generated 15 distinct clusters that parsed into 22 neuron types. These 22 neuron types were present across all three intestinal regions studied. Instead of attempting to flow sort neurons or neuronal nuclei from human intestine, the group relied on laser capture microdissection of histochemically stained myenteric ganglia collected from healthy young organ donors. The team used comparative analysis to remove genes expressed in surrounding muscle, producing a deep bulk data set specific to human myenteric ganglia. By comparing genes expressed in their human bulk RNA-seq data to those present in mouse myenteric neurons, the authors were able to identify genes that marked distinct types of enteric neurons in both species. The regional and cell type specific expression of these “marker genes” were assessed *in situ* via hybridization chain reaction to localize distinct neuron types in each bowel region for both species.^34,35^ Importantly, the May-Zhang study implemented a method for blocking lipofuscin auto-fluorescence that is problematic in visualization of human neurons and can confuse interpretation of labeling.^36^ The team’s analysis focused on localization of intrinsic primary afferent neurons (IPANs) *in situ* based on markers of somatostatin and calbindin-2 in parallel with Kelch Like Family Member 1 (*KLHL1*) present in murine IPANs. The team found that there is incomplete congruence of expression for genes that mark neuron types in mice compared to those expressed in human enteric neurons. Moreover, these authors determined that multiple genes exhibited regional differences in gene expression between the duodenum, ileum, and colon of mice, and most of these regional expression patterns were not detected in human tissues. These findings illustrate some cross species congruence, yet point to species distinctions, indicating the need for caution when extrapolating studies in mice for comparisons with human ENS studies.

Complementary ENS profiling from Morarach et al., in 2021 produced transcriptional signatures from enteric populations in small intestines of mice as fetal progenitors differentiated towards neuronal fates.^10^ Morarach and colleagues generated scRNA-seq profiles from neural crest-derived enteric neurons and glia based on *Wnt1-Cre:R26R-Tomato* expression with sequencing of 3260 cells at 15.5 dpc and 2733 cells at 18.5 dpc. To assess murine juvenile enteric neuronal diversity, the team utilized a *Baf53b-Cre* transgene that labels large numbers of enteric neurons to gain transcriptional profiles of an additional 4892 cells from the myenteric plexus of the small intestine at P21. From these data, the authors defined 12 main transcriptional enteric neuron cell states that mostly express either *Etv1* or *Bnc2* in a dichotomous manner. The authors rigorously validated all detected neuron types via immunohistochemistry for marker genes in each cluster. Their validation studies showed that two neuron subsets exhibit morphological features consistent with known morphology of IPANs. In their fetal datasets, the authors found the same *Etv1*, *Bnc2* transcriptional dichotomy in two “branch” trajectories of emerging enteric neurons over developmental time. This distinctive binary split in development of enteric neuron lineages contrasts with neurogenesis programs in the central nervous system and indicates that neuronal diversity in the ENS is elaborated post-mitotically after these initial branching events. From their transcriptional profiling of developing ENS, Morarach and colleagues noted expression of *Pbx3*, a transcription factor expressed at a transition point bordering two neuron clusters in their scRNA-seq data. They further showed that loss of this gene produced altered ratios of Calbindin+ neurons. The team’s work illustrates how careful attention to emerging transcriptional programs can identify key regulators that establish normal neuronal diversity in development.

Later, a study from Wright et al., 2021 produced snRNA-seq data for mouse enteric neurons and glia from young adult mouse intestine based on labeling with *Wnt1-CreERT2^Cre/WT^:R26R-H2BmCherry*+ crosses.^11^ The team analyzed numerous mouse reporter lines to identify this combination that produced bright labeling of ENS nuclei and avoided erroneous flow sort gating due to adherence of neurites and cellular fragments attached to negative cells, which can occur with cytoplasmic tdTomato labeling. The analysis identified seven distinct neuronal clusters from 635 adult neurons sampled. Wright and colleagues further profiled fetal cholinergic (*ChAT-EGFP-L10AD*+) and nitrergic (*Nos1-CreERT2^Cre/WT^;R26R-TdTomatoD*+) neurons producing scRNA-seq data for 707 neurons that distributed into 8 distinct clusters at 17.5 dpc. The authors noted multiple differentially expressed genes between neuron types were known regulatory factors and utilized conditional gene deletion strategies to assess whether deletion of individual genes altered abundance of myenteric neuron types. The team’s analysis demonstrated that *Tbx3* is required in the fetal ENS for normal abundance of Nos+ neurons. In contrast, deletion of other transcription factors (*Casz1*, *Tbx2*, and *Rbfox1*) from developing ENS did not alter abundance of cholinergic neurons. These studies suggest that production of some neuron subtypes may rely on more than one regulatory factor, a key finding for future directed differentiation of enteric neuron classes.

More recently, Guyer and colleagues in 2023 produced single cell multiome sequencing data from 17690 enteric glia from *Plp1::GFP* mice at P14.^13^ In addition, they also produced single cell RNA-seq data and single cell multiome sequencing for *Plp1::GFP* neurospheres derived from 12-14 week old mice. Using these data paired with immunofluorescence and RNAscope imaging, the authors confirm that enteric glial cells within ganglia are poised for undergoing neurogenesis in postnatal and adult stages.

Recently, Vincent et al. in 2023 produced a developmental ENS dataset comprised of *Ret^CFP/+^* and *Ret^CFP/CFP^* (loss of *Ret*) mutant cells.^14^ They utilized this dataset to probe how the ENS develops in the absence of *Ret* expression, which they found affects the inhibitory neuron subtype differentiation, the timing required for proper neuronal and glial fate decisions, and ENS cell cycle dynamics.

Also in 2023, Kulkarni et al. published as version of record in *eLife* scRNA-seq data from isolated longitudinal muscle-myenteric plexus (LM-MP).^15^ By mining this data set the authors argue for identification of a subset of adult enteric neurons that are *Wnt1*-cre negative, meaning that they are not derived from the enteric neural crest cells. Kulkarni et al. conclude that these non-neural crest neurons increase in number as both mice and humans age. Furthermore, the team presents evidence that these non-neural crest enteric neurons express markers suggestive of a mesoderm-derived origin. These results were highly disputed by the reviewers, as the study did not use flow cytometry via their multiple mesoderm cre reporters to isolate these cells, opting for scRNA-seq of all cells from the LM-MP in mice at both 6 months of age and between P10-P30. This particular scRNA-seq data were not sequenced at a depth typical for scRNA-seq studies, which could cause these data to be dissimilar to other mouse enteric neuron single cell RNA-seq datasets from the same ages. Despite this caveat, the reviewers do acknowledge that there is a population of enteric neurons that originate from a lineage independent from the neural crest.

This year, Schneider et al. in 2024 performed scRNA-seq of enteric nervous system at P5 from *Tyrosinase-Cre Bap1^fl/fl^* mice showing that fetal deletion of *Bap1*, a chromatin modifier, caused a shift in enteric neuron subtypes, but this is not the case at postnatal stages.^16^ This reveals another gene that is required for proper subtype proportions, similar to *Sox10* as found by Musser et al., 2015.^3^

Most recently, Zhou et al., 2024 performed scRNA-seq on three gastrointestinal regions at five developmental stages to attempt to assay cells that represent the migrating wavefront of enteric neural crest-derived cells.^17^ From the stomach, small intestine, and colon at 10.5, 12.5, 14.5, 17.5 dpc, and P21, they assayed a total of 4741 ENS cells. The authors claim that they identify a set of genes and a cluster of cells that are specifically upregulated in the migrating wavefront. In addition to this, the authors perform spatial transcriptomics via Stereo-seq on the mouse gut segment containing the migrating wavefront at 12.5 dpc, from which they find enrichment of the upregulated genes in their wavefront scRNA-seq cluster to be enriched at the Stereo-seq wavefront.

Each study profiling ENS progenitors and mature enteric neurons applied distinct approaches both technological and bioinformatic to produce single cell or nucleus RNA-seq data. Variables included differing mouse lines, ages, tissue dissociation methods, techniques for encapsulation/library production, and depth of sequencing.^5–17^ These differences have led to variation in the number of cells sampled, cell types detected, and cluster-specific marker gene expression.^2,18,19^ A main point of discussion in the ENS field centers on reaching consensus regarding how many ENS cell types exist and what methods should be prioritized for classifying and naming them for consistency across the literature. Combining enteric neuron scRNA-seq datasets in a strategic manner is one means to gain greater understanding of diversity of enteric neurons. We make the case below for this strategy and illustrate the process, outlining both advantages and caveats. Our goal is to demonstrate how meta-analysis (the process of pooling independent datasets together into one larger aggregate focused on the same question) can yield greater consistency of interpretation and offer a resource of similarly processed datasets for future mining by the ENS community.

### Why combine? The case for a meta-atlas of ENS scRNA-seq data

In scRNA-seq experiments, sampling of each cell type within a tissue can be challenging, especially when those cells are rare. This is particularly the case for the ENS, where enteric neurons are embedded within the bowel wall, making up a low percentage of total cell numbers in the tissue. Low abundance is further complicated by difficulties in tissue dissociation and loss of neuronal processes due to shear forces during isolation. Even after successful cell isolation, differences in cell numbers profiled and the labeling strategy used to fluorescently tag enteric neurons for flow sort isolation contribute substantially to differences in the data generated. Recent success in combining multiple independent scRNA-seq data sets into a single aggregate, or “meta-analyses”, illustrate how combinatorial approaches have been informative for studies of COVID patients, liver homeostasis, and atherosclerotic tissues.^37–39^ Each of these studies leveraged similarities across individual datasets and the increased cell number resulting from the integration to ask questions pertaining to the cell types of interest. They also successfully derived a consensus of cell type classifications and found novel cell subtypes that were not previously characterized from the individual datasets alone. Likewise, we take advantage of the similarities between ENS scRNA-seq datasets to successfully compile a “meta-atlas” of enteric neurons that offers a deeper resource for discrimination between enteric neuron types. This approach (Figure 1) reveals both commonalities between datasets as well as some differences between neuron types that were not as evident until data aggregation was performed.

**Figure 1:**
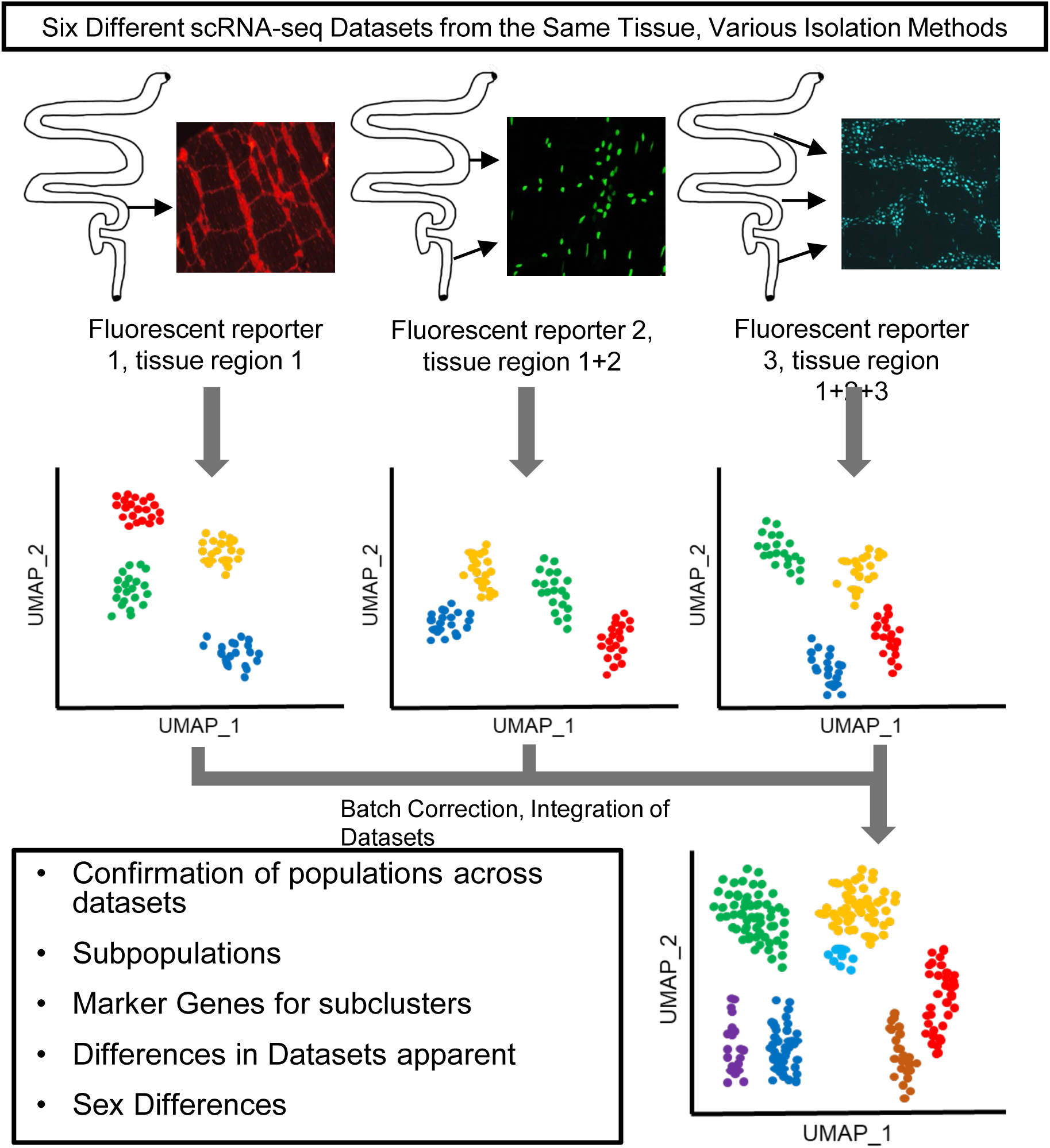
Overview of data integration process and analysis approach.

### Advantages of reprocessing scRNA-seq data

Comparably processed datasets are optimal for production of a meta-atlas. Typically, initial scRNA-seq data in its large FASTQ file format is aligned via tools like CellRanger that produce barcode, matrix, and gene files.^30^ These outputs are then imported into programs like Seurat for filtering, normalization, scaling, and dimensionality reduction to visualize differentially expressed genes.^21–25^ It is possible to obtain aligned files associated with each publication although these often include alterations that result from data processing. Some authors provide the completely processed Seurat object; however, these files are too large for easy sharing. Importantly, use of derivative Seurat objects for comparisons can be complicated by how each dataset was processed. Differences in quality control metrics or clustering parameters can cause cells to be dropped from further analysis, lead to retention of poor-quality cells, or cause cell clustering differences. Reprocessing aligned data, such as that found in raw counts matrices and using a consistent set of parameters avoids these complications and leads to greater confidence in the outcomes produced by cross-study comparisons.

To illustrate how a meta-atlas can be produced, we primarily relied on semi-processed data files (such as CellRanger outputs) from alignment to the mouse genome (version mm10). We focused our ENS meta-analysis on juvenile and adult ENS scRNA-seq datasets from the mouse, as these cell states are most alike (Table 1). In contrast, the developmental datasets, despite expressing pan-neuronal marker genes, contain immature progenitor populations that are less likely to align with mature neuron clusters and can exhibit transient expression of developmental genes that are not retained in mature neuron types. We sourced these data from either the Gene Expression Omnibus (GEO), the Single Cell Portal at the Broad Institute, or from mousebrain.org.^5,9,10,11,12^ Some of the Drokhlyansky datasets had been pre-processed to eliminate heterotypic cells (sequenced droplets that contain two or more cells or nuclei of different types) using MIRACL-seq algorithms. We relied on these processed datasets for convenience, as most groups interested in utilizing these datasets may not have the computational tools for genome alignment. While not perfect, this is a straightforward approach for many groups that do not have advanced bioinformatics capabilities.

### Combining datasets: process, gains, and caveats

#### Reprocessing each sc/snRNA-seq dataset

For data comparison and integration, we used the R package, Seurat, for quality control metrics and unsupervised clustering and Seurat’s SCTransform v2 to integrate each dataset’s sequencing replicates (referred to as runs).^21–25,27,28^ Initially, similarities between enteric neuron (EN) scRNA-seq datasets were assessed by several standard bioinformatic approaches. All data sets were plotted concurrently using the same uniform manifold approximation projection (UMAP) parameters (Figure 2A-F). UMAP plotting is a dimensionality reduction approach that shows how cells, represented as dots, are transcriptionally similar to each other as displayed by the distance between cells. Cells that are closer together are transcriptionally more similar than cells that are widely separated on a UMAP. These plots reveal the differences in cell numbers between datasets. Second, we evaluated expression of genes known to mark discrete enteric neuron types and displayed the expression of these marker genes for each data set using dot plots (Figure 2A’-F’). This approach revealed consistent detection of canonical neuron subtype markers across datasets, especially those with highest cell numbers. We also performed differential gene expression within each dataset to find putative cluster marker genes (Seurat’s FindAllMarkers), which can be found in Supplementary Tables 1-7.

**Figure 2:**
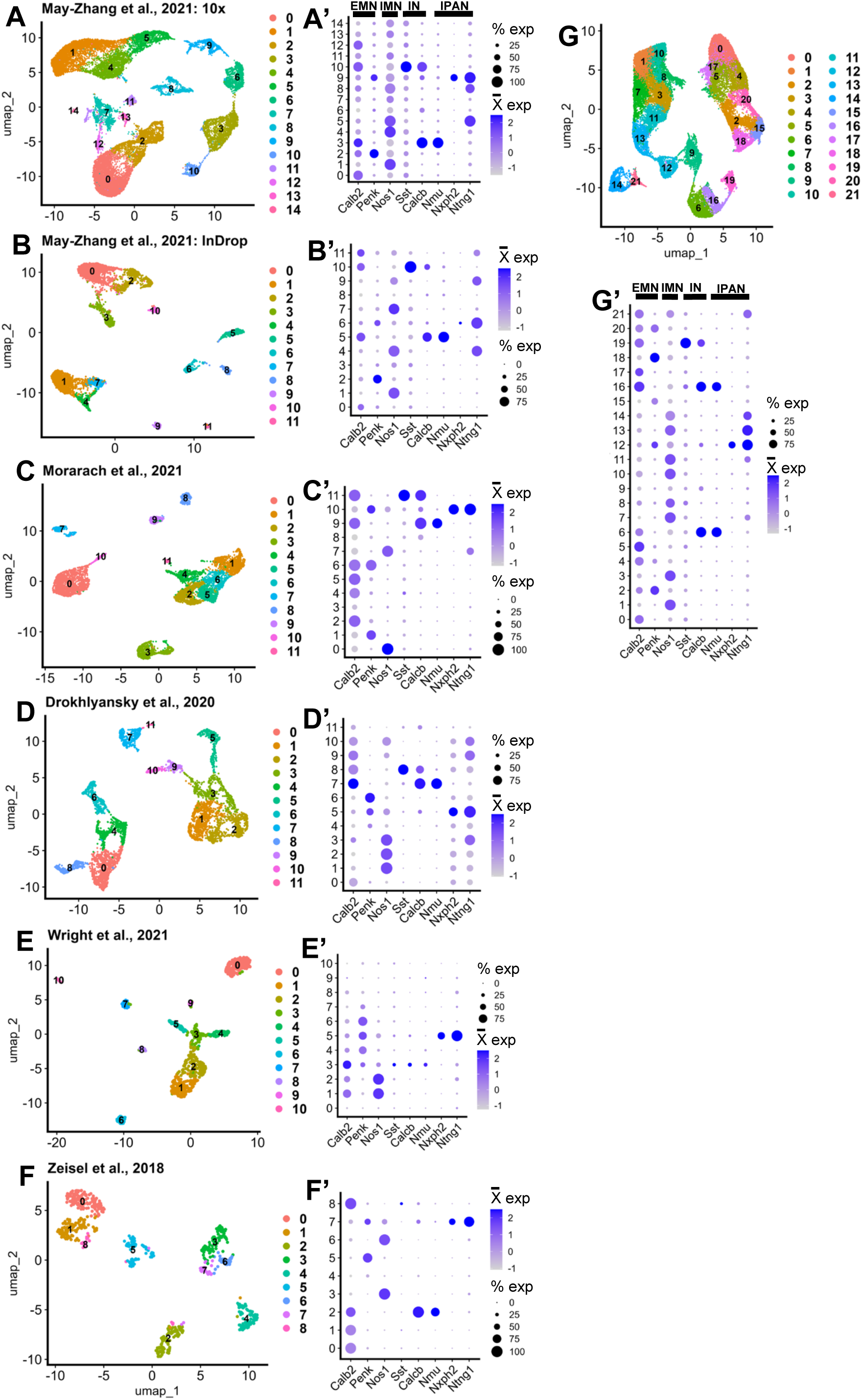
General consensus of enteric neuron types distributed across each single cell dataset. Panels **A**, **B**, **C**, **D**, **E, and F** show UMAP plots for each ENS dataset. Panels **A’**, **B’**, **C’**, **D’**, **E’, F’, and G’** display dot plots of established marker genes detected in enteric neuron clusters. Panel **G** shows the UMAP of the integrated ENS meta-atlas. EMN: excitatory motor neuron; IMN: inhibitory motor neuron; IN: interneuron; IPAN: intrinsic primary afferent neuron.

#### Removal of putative non-neuronal clusters pre-integration

We then focused our meta-analysis on clusters that were present in all datasets. In these initial comparisons, we noted that the May-Zhang 10X data contained several cell clusters that did not or mostly did not correspond to clusters in the other cohort datasets. Specifically, May-Zhang 10X reprocessed clusters 7, 11, 12, 13, and 14 (corresponding to clusters 4, 11, 12, 13, and 14 from May-Zhang et al., 2021’s in-publication processing) were not present or not detected in the final analyses of Drokhlyansky and Morarach.^5,9,10^ We estimated that May-Zhang 10X cluster 13 (in-publication cluster 12) appeared consistent with our reprocessing of Morarach et al., 2021’s cluster 3, which was marked as a mix of enteric glia and neurons in their Figure 1d (Figure 2C).^9,10^ These clusters express both glial and neuronal markers, and as a result were excluded from the original analysis by Morarach and colleagues.^10^ To specifically focus on differentiated neuron types, we excluded most of the clusters that were in May-Zhang et al., 2021 that expressed markers that were not shared across other datasets (Supplemental Figure 1A,B). We also excluded the cells from the Morarach dataset’s reprocessed cluster 3 for the same reasoning as the May-Zhang et al., 2021 unknown clusters (Supplemental Figure 1C). Retaining such dataset-specific clusters in the meta-atlas would not aid in determining cell identity for these clusters because the clusters do not gain additional cells when integrated with other datasets, and therefore do not have higher resolution. May-Zhang et al., 2021’s in-publication cluster 8 was an unassigned cell type, but expressed *Nos1*, *Gal*, and *Sst* (at lower levels), so we chose to leave corresponding reprocessed clusters with this expression pattern in the meta-atlas that did not express those non-neuronal genes.^9^ In a similar light, we also removed reprocessed cluster 0 in the Wright et al., 2021 dataset due to expression of glial cell markers (Supplemental Figure 1D). We postulate that the differences in cluster presence between datasets most likely has to do with the fluorescent reporters that were used to isolate the cells.

#### Integration of reprocessed datasets to generate a single cell transcriptomic meta-atlas of enteric neuron types

To integrate all datasets into one, we integrated by biological replicate, which resulted in the integrated dataset represented by a UMAP dimensionality reduction with 22 clusters generated via unsupervised clustering (Figure 2G). We confirmed that the characteristic neuron type marker genes initially detected in each individual dataset (Figure 2A’-F’) were consistently observed in the integrated dataset by re-examining these markers using dot plots (Figure 2G’). As expected, the markers are expressed in distinct patterns after integration. The process of batch correction is an analytic technique designed to remove effects in the data that are due to the methods by which the data are generated, or technical variation, allowing discernment of true similarities or differences between datasets upon integration. Batch correction can be performed by any of several approaches.^40^ In our processing we applied batch correction via SCTransform v2 integration.^27,28^ To demonstrate this batch correction approach performed appropriately, we show UMAP and PCA plots with cells and split by cluster (Figure 3A, Supplemental Figure 1E-G). If batch correction were not successful, the plots could show one of two possibilities. Either the cells from each dataset would not be dispersed relatively evenly throughout each cluster, or clusters of differing cell types would be merged.

**Figure 3:**
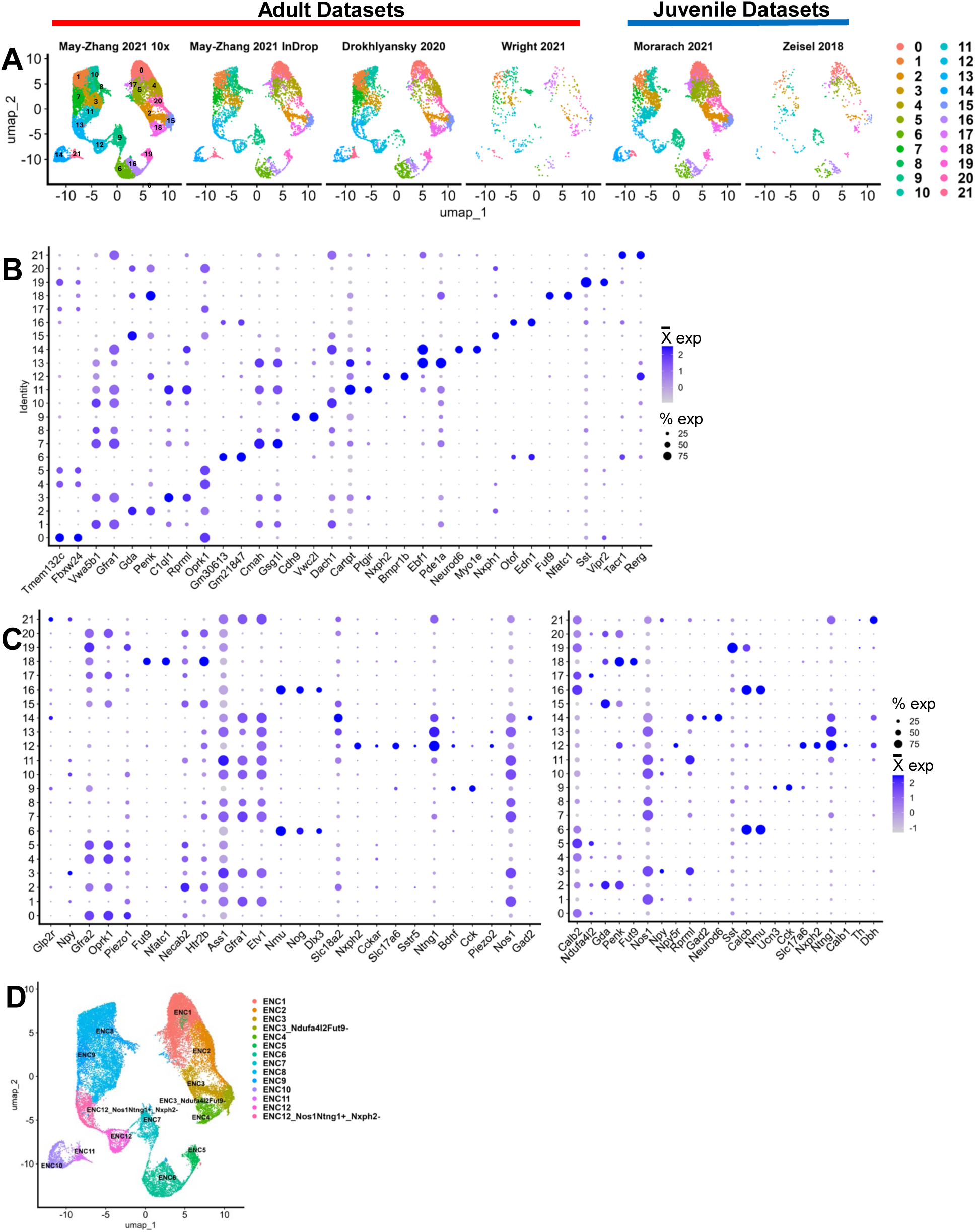
Post-integration gene expression in unsupervised clusters reveal distinct neuronal classes in relation to prior classifications. **A** UMAP from Figure 2G grouped and split by the dataset of origin, showing the results of batch correction of the cells from each dataset. **B** Dot plot of significant differentially expressed upregulated genes per cluster based on high log2 fold change and exclusive cluster expression. **C** Dotplot showing expression of marker genes from Dharshika & Gulbransen Figure 2 (Left) and Morarach and colleagues’ Figure 1j by cluster upon which the table classifications are based.^6,15^ **D** UMAP from Figure 2G grouped by putative “ENC” classifiers from Morarach et al., 2021.^6^

When combining datasets, we can achieve a depth of cell types that is greater than the individual datasets alone. For example, in our reprocessing of each individual dataset, each displayed varying numbers of neuronal types, and the datasets with the fewest cells had less diversity of neuronal types in their clustering. Interneurons and IPANs all clustered together in our reprocessing of the data from Wright et al., 2021 (cluster 5), and interneurons marked by *Sst* and IPANs marked by *Nxph2* and *Ntng1* were low in number in Zeisel et al., 2018 (clusters 8, 7; Figures 2E’,F’).^11,12^ In the meta-atlas dataset, these cells clustered with their appropriate counterparts from datasets with larger cell counts (Figures 2G,G’,3A).

#### Expression markers of meta-atlas neuron types detected via differential gene expression

With higher cell counts brought together by combining individual scRNA-seq datasets, one can further mine the meta-atlas for hypothesis generation and compare the integration to its original dataset components. We assessed a few basic questions to probe the integrated meta-atlas dataset. First, we aimed to identify genes that are significantly differentially upregulated in each cluster versus all other clusters, as these might assist in detection of expression markers of novel neuron types that were missed in the individual analyses. To accomplish this, we performed differential gene expression analysis for each cluster versus all others using Seurat’s FindAllMarkers function.^21–25^ We identified the top differentially upregulated genes per cluster by filtering the FindAllMarkers results to those genes that had average log2 fold change of greater than 1 and a significant Bonferroni adjusted p-value (p < 0.05; Supplemental Table 8). We then visually examined where on the UMAP these genes were expressed to prioritize those whose expression was primarily within a single neuron cluster. This process produced a list of putative marker genes for each of the resulting 22 clusters (Figure 3B).

#### Meta-atlas enteric neuron types defined by prior literature definitions

The meta-atlas can also allow us to compare proportions of cell types within each component dataset as well as use previous definitions of enteric neuron types to interrogate which clusters correspond to which neuron type. We have presented these data in table format and an accompanying dot plot (Table 2; Figure 3C). We use and add to the neuronal cell definitions from Dharshika & Gulbransen’s Figure 2, which shows the putative anatomy and morphology of cell types marked by gene expression from scRNA-seq, as well as use the “ENC” definitions from Morarach et al., 2021.^2,10^ Using their marker gene definitions, we plotted their expression for each cluster, showing distinct patterns (Figure 3C; Dharshika & Gulbransen: left dot plot; Morarach “ENCs”: right dot plot). Our clusters often expressed genes corresponding to conflicting cell types, so we opted to list those possibilities in Table 2. Of note, Dharshika & Gulbransen use *Slc18a2* expression as a marker for enteric glia.^2^ However, we found that this gene is also expressed in our neuronal cluster 14 (as well as lowly expressed in other clusters), which we posit is a secretomotor, vasodilator, inhibitory motor, or descending interneuron because of its expression of *Glp2r*, *Gfra1*, *Etv1*, and *Gad2*.^2^ Therefore, *Slc18a2* may not be a reliable marker exclusively for enteric glia *in situ*. From Table 2 and Figure 3C, we hypothesize that there are at least 15 shared distinct types of enteric neurons across these datasets, and that all the “ENCs” are represented here, with seemingly higher complexity.

**Table 2:** Distribution of cells across unsupervised clusters per dataset.

|  |  |  | Cell Counts Per Dataset |  |  |  |  |  |  |
| --- | --- | --- | --- | --- | --- | --- | --- | --- | --- |
| Putative Neuron Type | Putative “ENC” | Meta-Atlas Cluster | Adult |  |  |  | Juvenile |  | Meta-Atlas Total |
|  |  |  | May-Zhang 10X | May-Zhang InDrop | Drokhlyansky | Wright | Morarach | Zeisel |  |
| Excitatory Motor | ENC1 | 0 | 2341 | 557 | 401 | 6 | 628 | 97 | 4030 |
| Inhibitory Motor | ENC8 | 1 | 1653 | 334 | 562 | 59 | 190 | 24 | 2822 |
| Necab2+ Excitatory Motor | ENC3 | 2 | 917 | 272 | 231 | 34 | 771 | 31 | 2256 |
| Npy+ Inhibitory Motor | ENC8 | 3 | 1050 | 277 | 264 | 56 | 450 | 28 | 2125 |
| Excitatory Motor | ENC2 | 4 | 1187 | 239 | 313 | 16 | 301 | 56 | 2112 |
| Necab2+ Excitatory Motor | ENC1 | 5 | 946 | 289 | 100 | 7 | 688 | 52 | 2082 |
| Intrinsic Primary Afferent | ENC6 | 6 | 1187 | 195 | 213 | 8 | 120 | 45 | 1768 |
| Inhibitory Motor | ENC9 | 7 | 1039 | 214 | 342 | 37 | 115 | 5 | 1752 |
| Inhibitory Motor | ENC9 | 8 | 970 | 377 | 143 | 111 | 130 | 7 | 1738 |
| Piezo2-IPAN/Intestinofugal Afferent | ENC7 | 9 | 1075 | 201 | 60 | 10 | 270 | 79 | 1695 |
| Npy+ Inhibitory Motor | ENC8 | 10 | 905 | 180 | 313 | 44 | 225 | 17 | 1684 |
| Ntng1+ Inhibitory Motor | ENC8 | 11 | 819 | 143 | 145 | 25 | 415 | 37 | 1584 |
| IPAN/Interneuron | ENC12 | 12 | 817 | 209 | 310 | 50 | 140 | 21 | 1547 |
| Ntng1+ Inhibitory Motor | ENC12, Nxph2- | 13 | 986 | 197 | 220 | 12 | 99 | 8 | 1522 |
| Vip-, Glp2r+, Gad2+, Slc18a2+, Etv1+ Neurons (Secretomotor/Vasodilator, Inhibitory Motor, Descending Interneuron) | ENC10 | 14 | 859 | 166 | 159 | 9 | 234 | 10 | 1437 |
| Excitatory Motor | ENC3, Ndufa4l2, Fut9- | 15 | 526 | 144 | 253 | 18 | 223 | 17 | 1181 |
| Intrinsic Primary Afferent | ENC6 | 16 | 725 | 97 | 146 | 14 | 116 | 52 | 1150 |
| Necab2, Htr2b Downregulated, Excitatory Motor | ENC1 | 17 | 261 | 175 | 71 | 60 | 531 | 40 | 1138 |
| Excitatory Motor, Rare Variant | ENC4 | 18 | 482 | 127 | 264 | 42 | 135 | 26 | 1076 |
| Sst+, Oprk1- Excitatory-like Interneuron | ENC5 | 19 | 545 | 113 | 200 | 6 | 106 | 8 | 978 |
| Piezo1 Downregulated, Excitatory Motor | ENC2 | 20 | 512 | 104 | 118 | 12 | 201 | 11 | 958 |
| Glp2r+, Npy+, Ntng1+, Inhibitory Motor | ENC11 | 21 | 243 | 77 | 159 | 13 | 49 | 3 | 544 |

Table of putative neuron cell counts grouped by cluster and dataset of origin. Putative neuron types are derived
from Dharshika & Gulbransen Figure 2.<sup>15</sup>

#### Differential abundance of putative enteric neuron types detected by age and intestinal segment

Integrating these datasets gives us an opportunity to identify distinctions between datasets and various annotations such as age and tissue section. To understand the differences in clustering distribution across age and tissue type (Figure 4A,B, Table 2), we performed Monte-Carlo permutation tests for each cluster (Figure 4C)^41^. We found that clusters 7, 13, 1, and 8 had significant proportional bias towards adult (May-Zhang et al., 2021, Drokhlyansky et al., 2020, and Wright et al., 2021) cells while clusters 5, 2, and 17 were proportionally biased towards juvenile cells (Morarach et al., 2021, Zeisel et al., 2018; Figure 4C, left). We also found that at least clusters 21, 8, and 7 had proportional bias towards colonic cells while clusters 5, 7, and 9 were proportionally biased towards small intestinal cells (Figure 4C, right). These patterns could hint at how enteric neuron populations change over time and throughout the length of the intestine.

**Figure 4:**
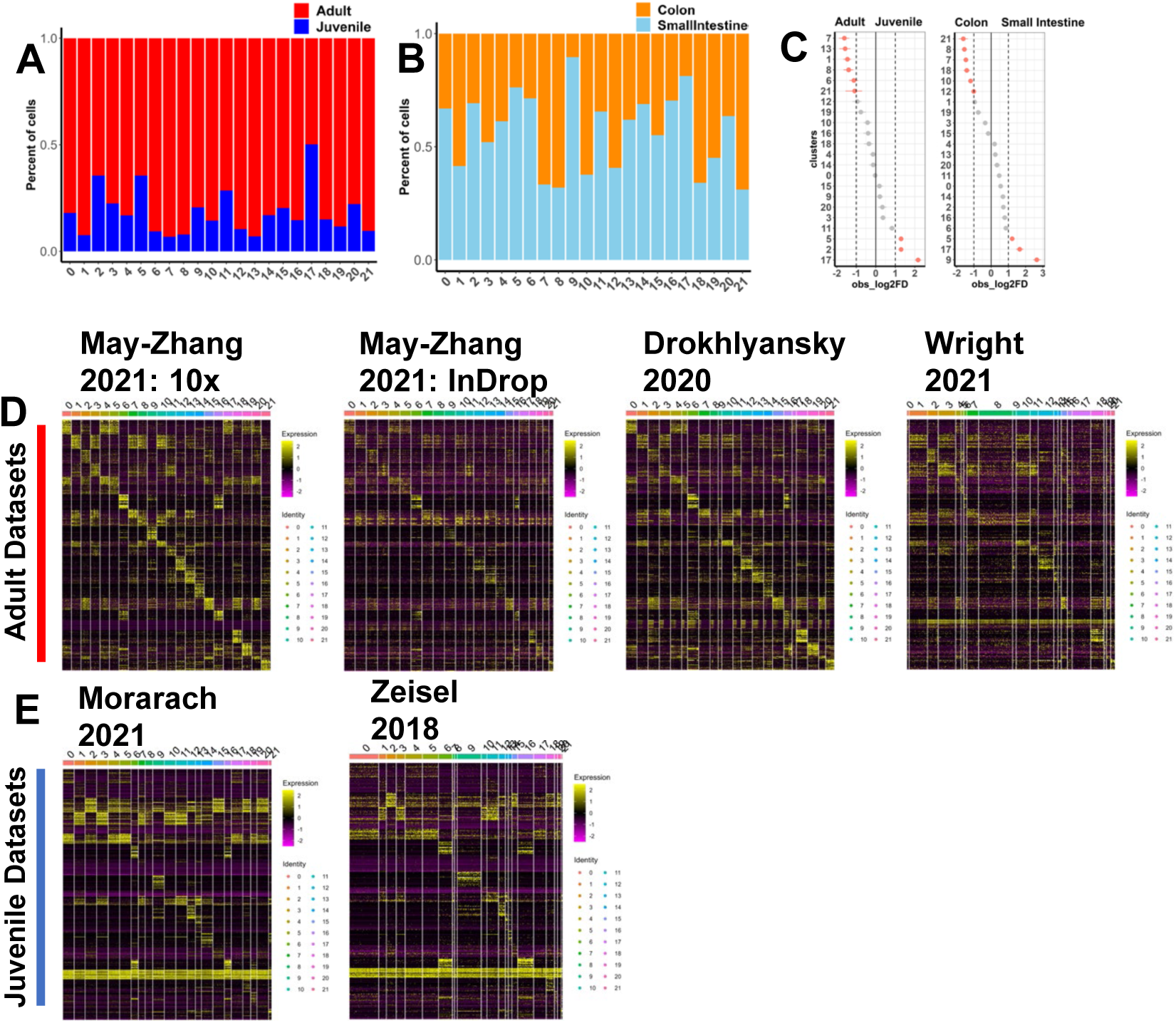
Post-integration cell distribution in unsupervised clusters, marker genes unveils distinct similarities and differences between datasets. **A** Stacked bar plot displays the distribution of adult and juvenile cells per cluster. **B** Stacked bar plot displays the distribution of colon and small intestine cells per cluster. **C** Permutation analysis displayed on a forest plot comparing the abundance of adult and juvenile cells per cluster and abundance of colonic and small intestine cells per cluster. **D** Heatmap displaying expression for the top 30 genes for each cluster from FindAllMarkers differential expression results of the meta-atlas dataset ranked by lowest p-value and highest log2 fold change plotted on the four adult datasets. **E** Heatmaps displaying the same 30 genes in **F** plotted on the two juvenile datasets.

#### Distinct expression patterns of top neuronal marker genes detected by age in the meta-atlas

To identify differences in expression between datasets, we used our top 30 genes per cluster from the FindAllMarkers results of the integrated dataset to plot heatmaps displaying expression patterns (Figure 4D,E, Supplementary Table 8). When we plot the gene expression for the subset adult datasets, the patterns of markers per cluster are relatively maintained, with dataset from Wright et al., 2021 displaying the worst congruence (Figure 4D). However, when plotting the subset juvenile Morarach et al., 2021 and Zeisel et al., 2018 datasets, many clusters do not have their own patterns of gene expression (Figure 4E). This leads us to conclude that the adult datasets, especially May-Zhang et al., 2021 10X dataset, are biasing the FindAllMarkers differential gene expression data, and that any further conclusions about cell type diversity must come from either the adult or the juvenile, but not both unless explicitly comparing the two general timepoints or finding what is common across age. Because of this, we decided to reintegrate only the adult datasets and use these data for downstream analyses (Figure 5A). We also perform FindAllMarkers differential gene expression analysis on these adult integrated data (Supplemental Table 9).

**Figure 5:**
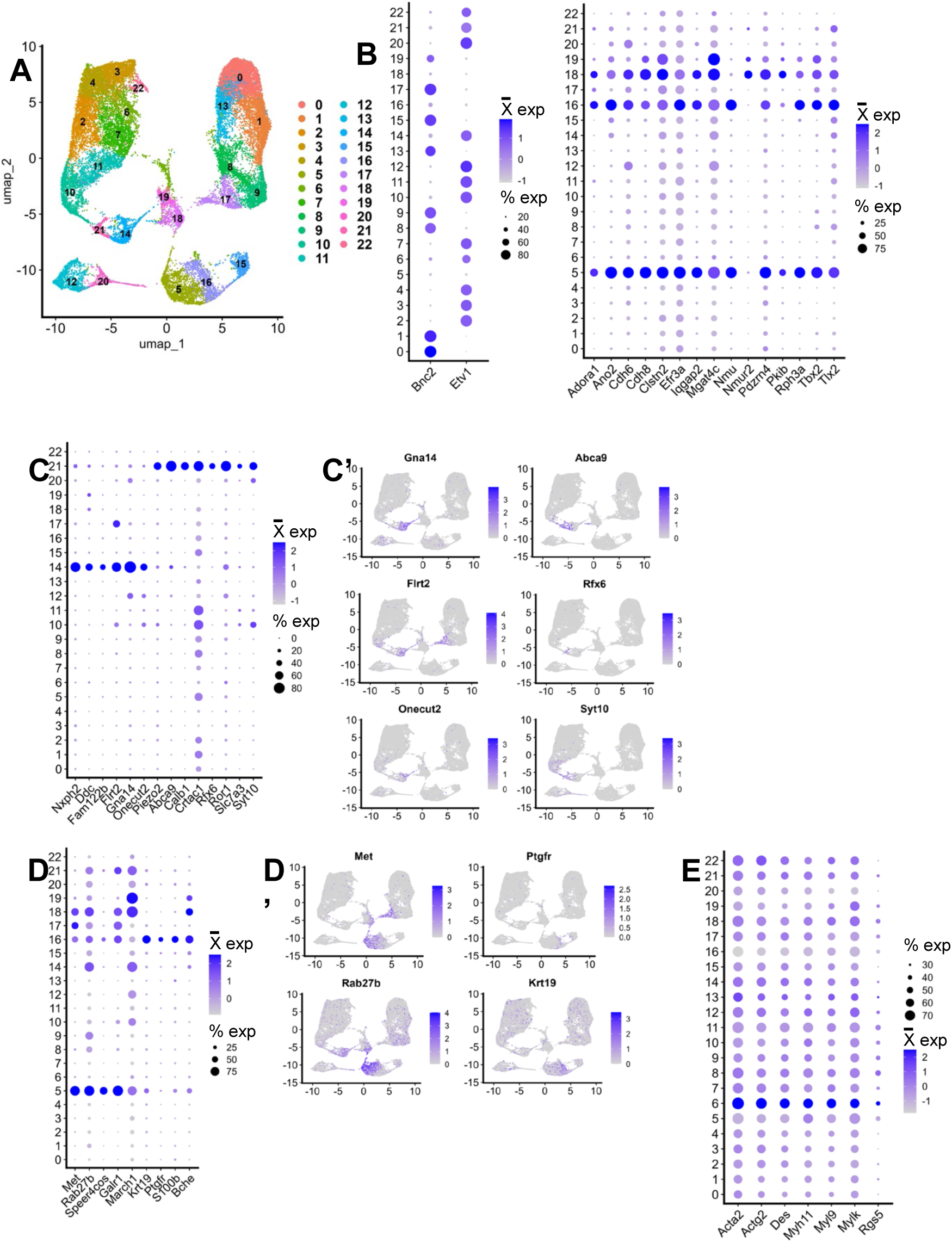
Probing the adult ENS meta-atlas for subclasses of enteric neurons. **A** UMAP of the new adult-only integrated meta-atlas. **B** Dot plots showing either Bnc2 and Etv1 expression is shown for all clusters (left) except 13, 18, and 19, which have distinct markers (right). **C** Dot plot showing expression of differentially expressed genes between clusters 14 and 21, with three representative genes’ expression per cluster plotted on the UMAPs in **C’**. **D** Dot plot showing expression of genes that typically mark non-neuronal types are expressed at low levels throughout the data but are most prominent in cluster 23. **E** Dot plot showing expression of differentially expressed genes between clusters 5 and 16, with two of these representative genes’ expression per cluster plotted on the UMAPs in **E’**.

#### Expression markers of clusters that do not express enteric neuron lineage markers Bnc2 and Etv1

With the effectively greater-per-cluster cell numbers, we were able to assess potential novel gene expression markers. For example, Morarach et al., 2021 found that the two “branched” neuronal lineages in the developing ENS were marked by the genes *Bnc2* and *Etv1,* respectively and the expression of these genes carried into juvenile stages. However, in their study (refer to Figures 2e,6f in Morarach et al., 2021), it appears there are cells that do not express *Bnc2* (ENC6, some of ENC7, end of Branch B) in the same way that the other branched linage expresses *Etv1* almost entirely.^10^ Following integration of the adult ENS single cell datasets, the distinct expression of *Bnc2* and *Etv1* is apparent (Figure 5B, left). This provides the opportunity to probe the adult meta-atlas at higher resolution than their original datasets to determine whether cells that lack *Bnc2* or *Etv1* express a gene or set of genes that distinguish these separate populations that might be expressed at fetal stages and maintained into adulthood. To explore this possibility, we took all our integrated clusters that expressed either *Bnc2* or *Etv1* and performed differential gene expression versus the clusters that do not appreciably express either of these two genes (Supplementary Table 10). At least 13 prominent genes emerged from this process (Figure 5B, right), each of which mark the clusters that do not express *Bnc2* or *Etv1*. For example, *Tbx2* was previously shown to have differential expression between neuronal types via immunohistochemistry and cells expressing this gene were enriched in cholinergic enteric neurons.^11^ Loss of this gene, however, did not alter density of *Chat*+ enteric neurons. Postnatal motility experiments in adult or juvenile mice were not possible because *Tbx2* mutants died shortly after birth.^11^ *Tlx2* has been established to interact with *Phox2a/b* in neural crest-derived cell development, with three *Tlx2* knockout mouse models having ENS defects^42–46^. *Iqgap2*, seemingly the most specific marker, has been reported to be required for inflammatory responses in the mouse colon.^47^ Identification of these genes as markers that distinguish these neurons from those expressing *Bnc2*/*Etv1* offers the opportunity to interrogate these cell types further in adult and younger stages.

**Figure 6:**
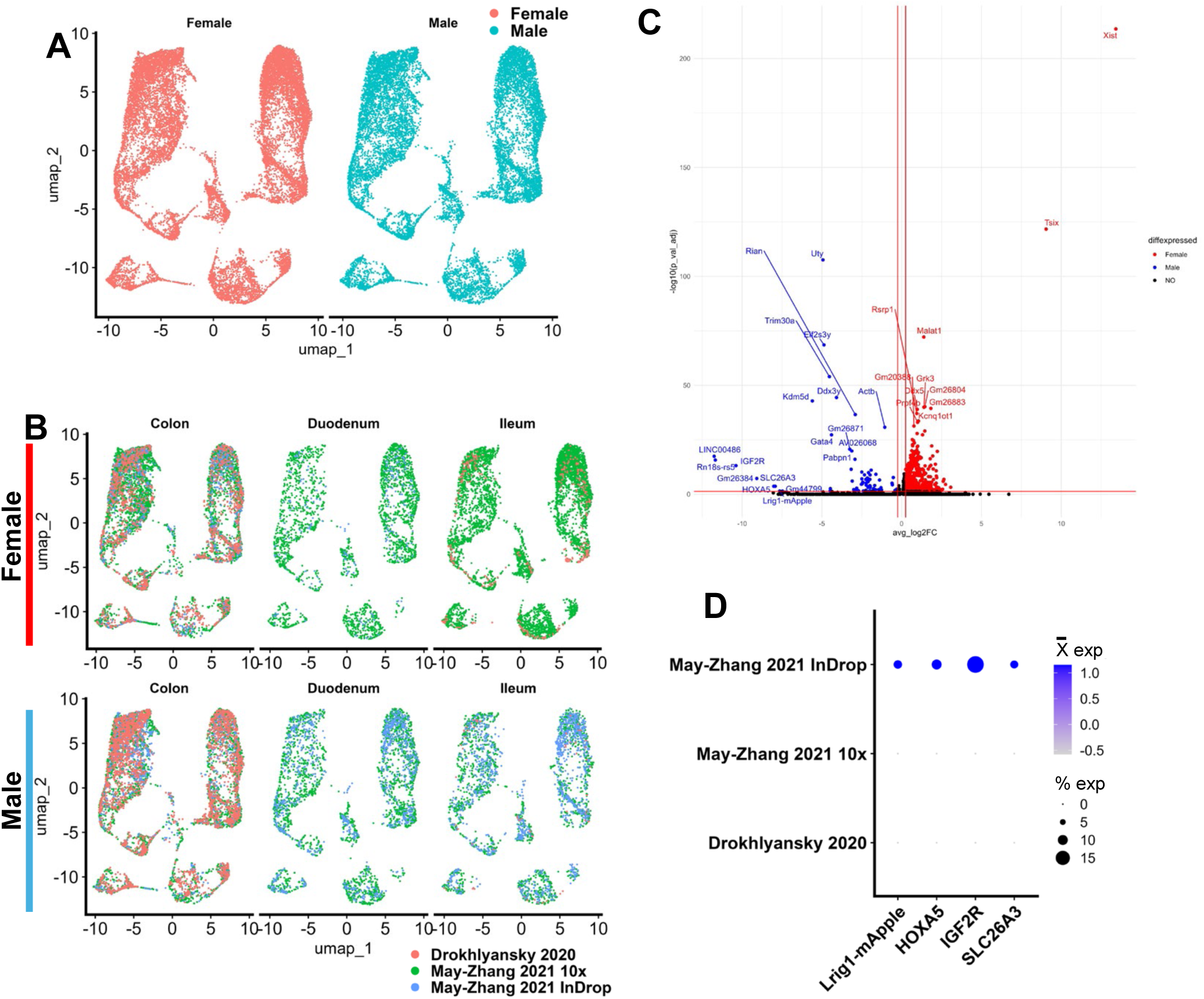
Sex and dataset differences identified in the adult meta-atlas. **A** Distribution of sex in the dataset visualized through a split UMAP. **B** UMAP displaying distribution of datasets of origin in the adult meta-atlas split by intestinal segment and stacked by sex. **C** Volcano plotting differential gene expression in cluster 6 between male and female cells showing erroneous genes upregulated in males. **D** Expression dot plot showing expression of “human” and a fluorescent fusion protein gene across datasets of origin.

#### Differential gene expression analysis identifies differences between Nxph2+ ENC12 and Nmu+ ENC6 subclusters

Increasing cell numbers through data integration also allows further subdivision of clusters to resolve previously unappreciated neuron subtypes. Morarach et al., 2021 reported that their cluster ENC12 was complex in both its scRNA-seq profile and in morphology observed by immunohistochemistry. Subtypes of this cluster were evident in their analysis, although cell numbers were fewer than other clusters in their dataset. We probed the adult integrated meta-atlas dataset to determine whether any of these subtypes could be further discerned or characterized within the adult stages. In the adult meta-atlas, clusters 14 and 21 coincided with Morarach ENC12 as verified by expression of ENC12 marker gene *Nxph2* and ENC12 subtype marker *Piezo2*, respectively (Figure 5C). Utilizing the higher adult meta-atlas cell count for ENC12 and Seurat’s FindMarkers differential gene expression analysis, we found additional adult marker genes for these subclusters, including *Abca9* and *Syt10* for cluster 21 and *Flrt2* and *Gna14* for cluster 14 (Supplementary Table 11, Figure 5C,C’).

When performing unsupervised clustering in both the complete and adult-only meta-atlas, we noticed that the *Nmu*+ cluster was split into two subclusters (Figures 2G, 3A, 5A). To determine whether this clustering could be reflective of biology and two enteric neuronal subtypes, we performed differential gene expression between the adult meta-atlas clusters 5 and 16 (Supplementary Table 12). By visual inspection of the top differentially expressed genes per cluster for specificity, we found at least 8 differentially expressed genes that may be tested for functional relevance (Figure 5D,D’). Marking cluster 5, we find hepatocyte growth factor (HGF) receptor *Met* and previously identified *Rab27b* (Figure 5D,D’).^9^ Marking cluster 16 we find *Ptgfr* and keratin gene *Krt19* (Figure 5D,D’).

#### Non-neuronal expression markers persist in the adult meta-atlas

When integrating datasets from different sources, clusters that might not have been previously identified could manifest. In the complete integrated and adult meta-atlas datasets, we observed that meta-atlas cluster 8 and adult meta-atlas cluster 6 express genes that typically mark non-neuronal cell types (*Acta2*, *Actg2*, *Des*, *Myh11*, *Myl9*, *Mylk*, *Rgs5*; Figures 2G, 5E, Supplemental Figure 2A). When split by dataset of origin, the adult integrated dataset UMAP shows that most of these cells come from the May-Zhang et al., 2021 datasets (Supplementary Figure 2B). When these non-neuronal marker genes are mapped back to the original May-Zhang et al., 2021 datasets, we find that they are widely expressed throughout, with highest expression in their unclassified clusters (Figure 2A,B, Supplementary Figure 2C,D). These genes were also expressed in the other two datasets. In the reprocessed Morarach dataset, there is limited expression of these non-neuronal genes in cluster 3 (Figure 2C, Supplementary Figure 2E). Expression is more dispersed in the reprocessed Drokhlyansky dataset, with highest expression in clusters 1, 6, and 7 (Figure 2D, Supplementary Figure 2F). It’s possible this dispersed expression of non-neuronal genes could be the result of ambient RNA due to muscle cell lysis during generation of nuclei from laminar muscle preparations. This expression could also arise from hetero-doublets (two nuclei of distinct cell types in the same droplet), although this is less likely given the use of FACS-sorting with size gating. However, Zeisel and colleagues reported the presence of enteric mesothelial fibroblasts marked by expression of *Pth1r*, *Kcnj8*, *Abcc9*, *Cd82*, *Tagln*, *Dcn*, *Lum*, *Pdgfra*, *Sox10*, *Aldh1a3*, and *Anxa11*.^12^ In the integrated adult meta-atlas, a subset of these genes (*Kcnj8*, *Abcc9*, *Cd82*, *Tagln*, *Dcn*, *Lum*, and *Pdgfra*) exhibit expression mostly in cluster 6 (Figure 5A, Supplementary Figure 2G). Whether the differences between datasets are due to technical challenges in sample processing or reflect biologically relevant cell types or states remains to be determined.

#### Sex differences validated and major dataset differences identified through differential gene expression

Most of the reports used to source data for the integrations of this analysis used recorded or estimated sex to either display cell distribution by sex, account for sex in differential gene expression (DGE) analysis, or perform DGE analysis by this recorded or estimated sex metadata. Because the aggregate higher cell quantities exceeded those of each individual dataset, we decided to use the opportunity to probe the meta-atlas for sex differences via DGE in our adult integrated data. However, since Wright et al., 2021 did not record sex, and since the contributions to the overall data are smaller than the other three, we elected to remove the Wright et al scRNA-seq runs for the sex-differences analysis.^11^ In order to track sex, we used both the annotations from the Drokhlyansky et al., 2020 and May-Zhang et al., 2021 publications, as well as the expression of known genes *Ddx3y*, *Uty*, *Tsix*, and *Xist* (Supplemental Figure 3A,B,D,E,G).^5,9^ Before going forward with the DGE analysis, we looked at the proportions of each dataset’s cells per sex per tissue segment. This approach revealed an obvious bias in the small intestine segments for the data in May-Zhang et al., 2021’s 10X data in females (8164 May-Zhang 10X neurons versus 641 other dataset neurons) and both the May-Zhang 10X and InDrop datasets in males (3638 and 2227 May-Zhang 2021 10X and InDrop neurons, respectively, versus 63 Drokhlyansky 2020 neurons; Figures 6A,B). These inequalities in initial input cell numbers may skew the analysis and result in differentially expressed genes that can be explained by the dataset of origin instead of sex. In spite of this limitation, we performed DGE by sex per cluster. This revealed an unexpected and concerning issue in the data: the presence of “human” gene symbols and fluorescent protein fusion-coding genes such as *Lrig1-mApple* were differentially expressed and upregulated in mostly males (cluster 6, Figure 6C). These genes should not be present in these data and indicate that there was some error in the alignment process during initial processing of one of these datasets. To determine which dataset was the source of these genes, we plotted a select few of these problematic genes by dataset, which revealed that the InDrop data from May-Zhang et al., 2021 expressed these exclusively (Figure 6D). In their publication, May-Zhang et al. were unable to include the InDrop data in the final analysis because they did not integrate well with their 10X datasets. However, they integrated well here, possibly due to the more recent software packages used (Seurat V2 to V5) or the inclusion of more datasets (Figures 2G,3A,5A, Supplementary Figure 1E-G).^9^ Going back to review which alignment software was used for the InDrop runs in May-Zhang et al., 2021, we found these alignments relied on the DropEST pipeline in contrast to the 10X Genomics datasets that used CellRanger.^5,9,30^ We presume that the use of a different mapping reference with the DropEST pipeline is the origin of the unexpected gene symbols. To retain the InDrop run data for the sex analysis, we first removed any obvious erroneous genes from the integrated dataset. We then performed DGE analysis with the InDrop data compared against the 10X Genomics assayed cells in the integrated dataset to find the top genes that were enriched in the InDrop data, with the goal of identifying erroneously aligned genes that were not immediately obvious (Supplementary Table 13). In total, we removed 8764 obviously incorrect annotations from the integrated data due to sequence alignment to human genome regions or plasmid vectors and subsequently reran the DGE analysis by sex per cluster and for the data overall. Generally, we observed comparable top differentially expressed genes (ranked by Bonferroni-adjusted p-value, average Log_2_FoldChange, and absolute value of the percent difference between male and female cells that express a gene) for this cleaned analysis as obtained from the initial analysis that included the erroneous alignment features. In males, as expected, we see upregulation of *Uty*, *Kdm5d*, *Ddx3y*, and *Eif2s3y* and in females we see, as expected, upregulation of, in some clusters, *Cntnap5a*, *Xist* and *Tsix* (Supplementary Table 14). We also observe that *Malat1* is also differentially upregulated consistently across clusters in females (Supplementary Table 14). We then used logistic regression to regress out the effects of dataset of origin (i.e., May-Zhang 2021 10X, May-Zhang 2021 InDrop, Drokhlyansky 2020) while performing differential gene expression analysis by sex per cluster and per tissue segment (duodenum, ileum, colon; Supplementary Table 15). The strongest differentially expressed genes between male and female in each cluster for each intestinal region from this analysis were the same as previously stated. *Malat1* is differentially expressed in specific clusters in specific intestinal segments although always upregulated in female data (Supplementary Table 15).

## DISCUSSION

Here we have illustrated how the integration of six individual scRNA-seq or snRNA-seq datasets provides higher power to statistical analyses, while also adding more complexity. Meta-analysis of juvenile to adult mouse EN scRNA-seq datasets produced higher cell numbers for cell types shared across datasets, enabling verification of prior enteric neuron classifications, extending the identification of additional marker genes, and offering greater resolution for the subdivision of clusters. Our approach also allowed us to identify differences between datasets including major expression differences across age, differences in cell type distribution, and identification of clusters and cell populations that are specifically enriched within individual datasets or intestinal segments. In our permutation analyses, we found that a group of putative ENC12-like, *Ntng1*+ inhibitory motor (cluster 13), ENC8-like inhibitory motor (cluster 1), and ENC9-like, inhibitory motor (clusters 7, 8) neuron types were upregulated in adult, and that ENC1 and 3-like, *Necab2*+ excitatory motor (clusters 2, 5) and ENC1-like, *Necab* and *Htr2b* downregulated excitatory motor neuron types were upregulated in juvenile ages (Figures 3D,4C, Table 2). In addition, we found that putative inhibitory motor neurons (clusters 21, 8, 7, 10), a rare variant of excitatory motor (clusters 18), and an intrinsic primary afferent neuron or interneuron type (cluster 12) are upregulated in the colon. Finally, we found that inhibitory motor (clusters 5, 17) and a *Piezo2*-, ENC7-like neuronal types are upregulated in the small intestine. These findings indicate that the ENS may change in composition over time and throughout intestine segment, but this will require experimental validation. There was some overlap observed between clusters that significantly differ between age and intestine segment, which may indicate dataset-specific effects in the permutation analysis. Our attempt to identify sex-specific differential gene expression analysis confirmed previous findings. but it is difficult to know whether the novel differentially expressed genes, including *Malat1*, are being driven by sex or by a dataset-specific effect, even after regressing out dataset metadata.

In our analysis of the adult meta-atlas, we found gene expression markers of subclusters of both *Nxph2*+ “ENC12” and *Nmu*+ “ENC6”. In addition to previously identified *Piezo2* and *Nxph2*, we extended the known marker genes for “ENC12” subcluster 14, with differential expression of *Gna14*, *Flrt2*, and *Onecut2*, while subcluster 21 expresses *Abca9*, *Rfx6*, and *Syt10* when these subclusters are compared (Figures 5C,C’). *Flrt2* has novel tumor suppressor activity in breast cancer and localizes to pre- and post-synapses in the postnatal developing hippocampus where it may play a role in synapse formation.^48,49^ *Gna14* is differentially expressed in *Pou3f3*+ versus *Pou3f3*-immature enteric neurons at 17.5 dpc and was also reported as a marker gene of the entire ENC12 cluster by Morarach, now shown to be a marker of ENC12 “subcluster” cluster 14.^10,11^ *Abca9* was recently identified as a novel gene involved in triple-negative breast cancer.^50^ *Nmu*+ ENC6 previously was probed for subclusters in May-Zhang et al., 2021’s analysis across the intestinal segments (May-Zhang et al., 2021 Sup. Figure 5A).^9^ However, this cluster did not immediately split via unsupervised clustering of the entire dataset in May-Zhang et al., 2021 like it did in our adult meta-atlas. Using these unsupervised clusters as subclusters of *Nmu*+ ENC6, we found that subcluster 5 differentially expressed *Met* and *Rab27b* (Figure 5D,D’). Interestingly, marking *Nmu*+ cluster 5, as well as clusters 17 and 18, we found that the gene coding for the hepatocyte growth factor (HGF) receptor *Met*, which has been shown to be important for development of a subset of intrinsic primary afferent neurons (IPANs) which regulate motility and injury response pathways in the intestine, exhibited differential expression for cluster 5 (Figure 5D,D’).^51^ *Met* has recently been suggested as a marker of a putative mesoderm-derived enteric neuron lineage in the aged intestine. However, further validation using FACS to enrich these putative Met+ populations is needed.^15^ *Nmu*+ cluster 5 also differentially upregulated *Rab27b*, which was also found to be mostly restricted to a subset of *Nmu*+ neuronal cells in the integrated findings from May-Zhang et al., 2021 (May-Zhang et al., 2021 Sup. Fig. 5A; Figure 5D,D’).^9^ We also found that *Krt19*, a keratin gene, was differentially upregulated and expressed selectively in *Nmu*+ subcluster 16. We validated that expression of *Krt19* was also enriched in cluster ENT9 in the juvenile dataset from Zeisel et al., 2018 (not explicitly in the publication, but on MouseBrain.org), which is an *Nmu*+ cluster (Figure 5D,D’).^12,52^ Finally, *Ptgfr* was found to be differentially upregulated in *Nmu*+ cluster 16, and it is expressed mainly in a subset of this cluster (Figure 5D,D’). *Ptgfr* was found to be enriched in clusters ENC5 and ENC6 in the dataset from Morarach et al., 2021, but seems to be restricted to this *Nmu*+ subcluster in these later adult stages (Morarach et al., 2021 Figure 2a, left; Figure 5D,D’).^10^ Like the subclustering described in the previous section for “ENC12”, these findings must be validated with *in situ* experiments or flow sort purification to determine whether these initial findings reflect genuine enteric neuronal subtypes. Given the rarity of these neuron subclusters parsed by the additional marker genes, future validation efforts will need to incorporate significant enrichment strategies to derive experimental evidence that these subclusters reflect authentic enteric neuron subtypes within the adult ENC6 and ENC12.

The potential applications of an enteric neuron meta-atlas are tremendous, as this framework can be used to assess changes in cell profiles with age, disease, diet, or microbiome. Multiple groups have posted individual data sets online for continued access.^5–17^ However, to date no single repository exists for aggregated ENS data that is designed to facilitate access by investigators without bioinformatics expertise. To encourage further efforts and begin to address this challenge for the field, we offer the R objects and code from the present meta-atlas compilation on Open Science Framework so other investigators can readily extend and mine this meta-atlas going forward.

Our meta-atlas relied upon robust batch correction approaches to reduce variation between datasets as is typically done. However, recent advances in barcoding and multiplexing strategies position the field for simultaneous sequencing of distinct samples that will facilitate comparative analysis and further reduce variation.^53,54^ Multiplexing would be particularly useful for analysis of ENS change across the lifespan as iterative sampling for a single mouse line could be readily performed by a single laboratory. Challenges will remain for controlling variance between studies due to differences in transgenic lines, genetic background of strains, or diet. Reporting specifics of these variables in each publication is essential so that these factors can be accounted for and noted when differences are observed between datasets. Variation among human studies will remain a greater challenge compared to the ability to control environment, including microbiome, in rodents.

Choosing to process these data in readily available, previously aligned forms, has benefits and detriments in a meta-analysis such as this. For example, if one does not have access to enough computational resources to perform alignment via a program such as CellRanger, which requires at least 64GB of RAM, it is easy to download these processed data and see the results that may match the results from each publication. However, differences in alignment approaches, mapping references, or other data processing methods from each publication can lead to complications, such as was the case for the InDrop datasets from May-Zhang et al., 2021. Reprocessing and realigning the FASTQ files could potentially offer cleaner results across all datasets, due to greater uniformity of gene symbols used for combining the counts for each gene across all datasets during the merging process. Otherwise, gene synonyms and alternative aliases across different mapping references can prevent counts from being properly binned in the resulting matrix.

Importantly, the field still faces the challenge of profiling young, healthy adult <u>human</u> enteric neurons at single cell resolution. The prior May-Zhang study relied on laser capture to gather entire ganglia sections from young adults. Drokhlyansky and colleagues used MIRACL-seq to assess RNA from older colorectal cancer patients.^5^ Because enteric neurons are affected by colorectal cancer, isolation of single enteric neurons for transcriptional profiling from young healthy adults is still needed.^55,56^ While optimized methods of tissue dissociation that maintain neuronal viability would enable this process, use of frozen tissue isolates to generate nuclei, combined with sequencing of vast cell numbers as sequencing costs decline, will likely succeed in circumventing this issue.^57^

A greater challenge for the ENS field will be linking the emerging transcriptional profiles of cell types with the historical morphological and electrophysiological data that has previously been the gold standard for classifying enteric neurons. Expression of single immunohistochemical markers has in some cases facilitated linking a transcriptionally defined neuron cluster with prior knowledge of neuron classes.^10^ May-Zhang and colleagues relied upon expression of known markers detected within some neuron types to propose the neuronal identity of clusters in their single cell data.^9^ One approach could be to sequence single neurons after electrophysiological studies as has been done for brain neurons.^58^ However, culture of enteric neurons to evaluate their polarization profiles may alter gene expression patterns. As sequencing technologies advance, it is more likely that spatial sequencing or transcriptional profiling of live cells will lead to success in defining the transcriptome of healthy human enteric neurons in situ.^59^

We acknowledge that our approach lacks validation *in situ* or *in vivo* that will be essential to cofirm the cell subtypes revealed in this study reflect authentic enteric neuron subtypes. We encourage those in ENS neurobiology to further investigate these potential novel neuronal classes to gain further insights for the field.

Finally, while there is great excitement given the advances in technology for both producing and analyzing scRNA-seq, we acknowledge other processes beyond transcription are important for cell identity. Translation, post-translational modifications, and protein turnover may all be at work in producing the final identity and functionality of enteric neurons. Integration of transcriptional profiles, multiplex immunolabeling, proteomics, and lipidomics will aid in realizing the full diversity of neurons within the ENS.

## CONCLUSION

In this work, we performed a synthesis of publications assessing the enteric nervous system at a single cell level. We shared key insights from an integrated mouse meta-atlas of both adult and juvenile and adult single cell and nucleus RNA-seq enteric neurons. In our atlas, all previously annotated enteric neuronal types are present, and were further subtyped using known and newly identified marker genes. We also identified dataset differences, sex differences by cluster and by intestinal segment, and age differences in our enteric neuron meta-atlas. These findings have great potential to improve our understanding of the ENS, following validation by the ENS research community.

---

**Supplementary Figure 1:**
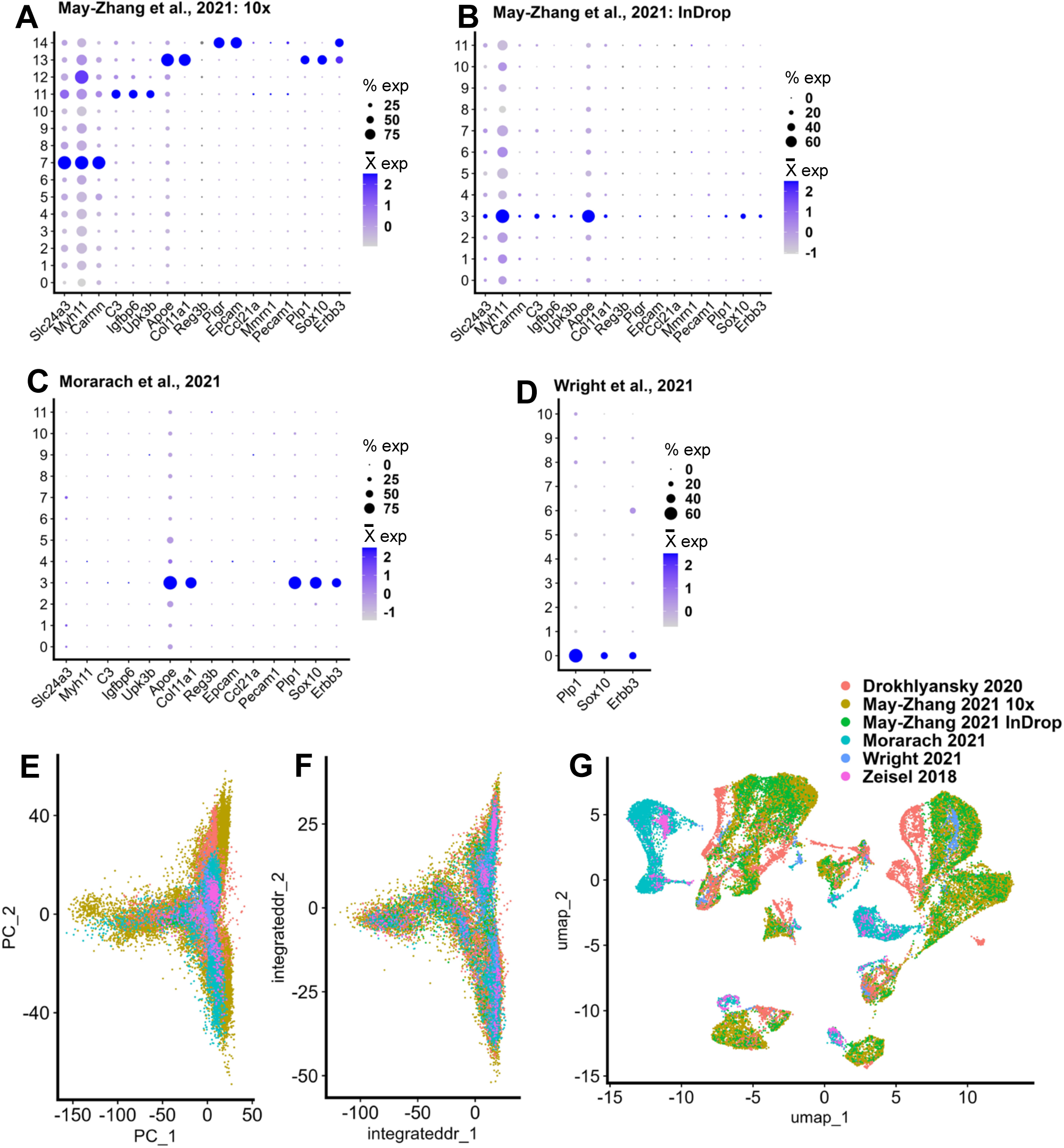
Identification of putative non-neuronal clusters to remove for effective meta-atlas generation and batch correction. **A-C** Dot plots displaying expression of gene markers of unknown clusters from May-Zhang et al., 2021 for both May-Zhang et al., 2021 datasets and the Morarach et al., 2021 juvenile dataset. **D** Expression of glial-like markers in the Wright et al., 2021 dataset on a dot plot. **E** Pre- and **F** post-integration PCA, showing proper integration of enteric neuron datasets. **G** Pre-integration UMAP of the enteric neuron meta-atlas.

**Supplementary Figure 2:**
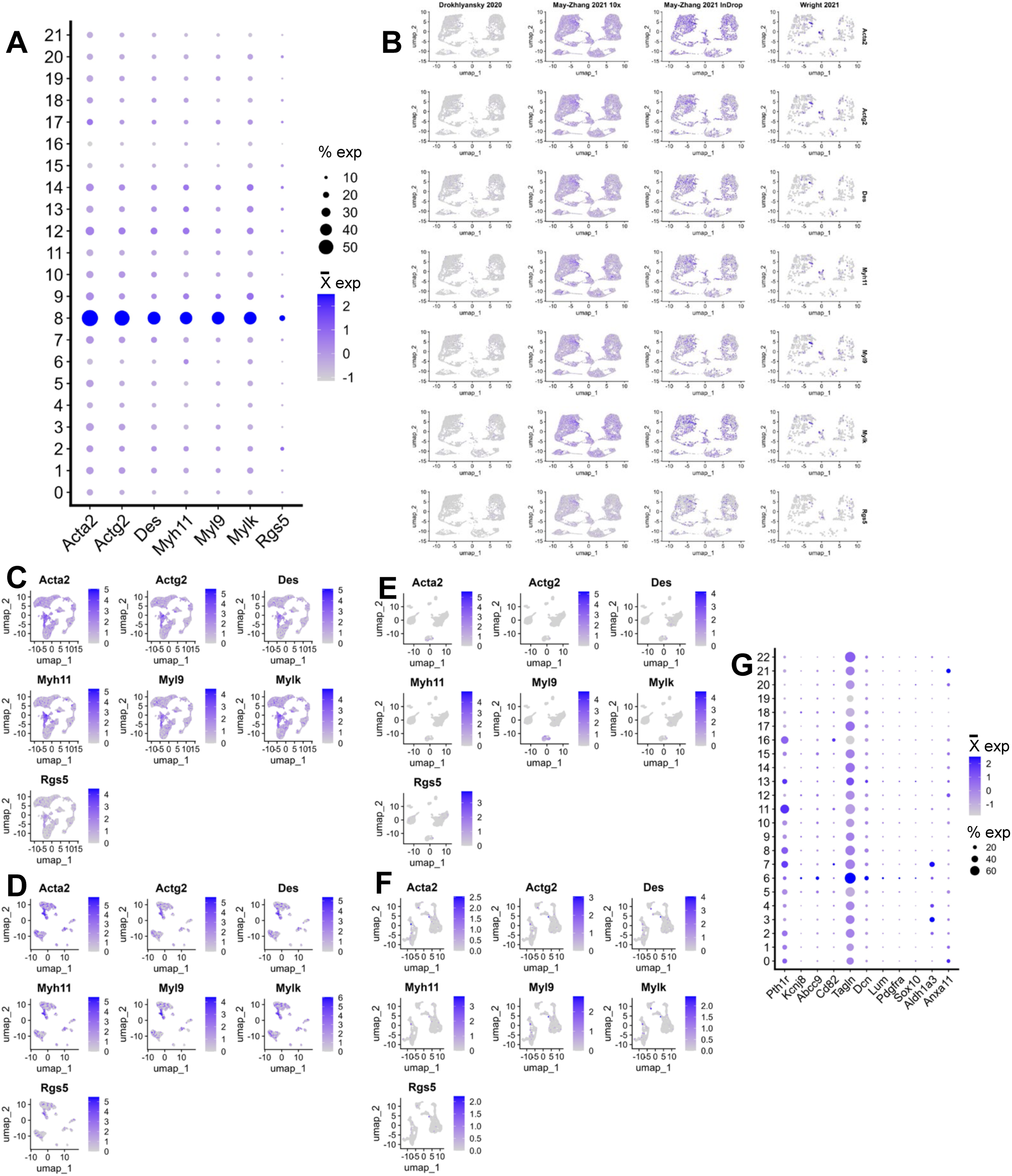
Muscle-like and enteric mesothelial fibroblast gene expression in the juvenile and adult meta-atlas. **A** Dot plot displaying expression of “muscle” expression markers in the combined juvenile and adult meta-atlas. **B** UMAPs showing expression of genes from **A** in the adult meta-atlas split by dataset, which identifies May-Zhang et al., 2021 as the main source of this gene expression. **C-F** UMAPs showing expression of genes from **A** in the May-Zhang et al., 2021 10X (**C**), InDrop (**D**), Morarach et al., 2021 (**E**), and Drokhlyansky et al., 2020 (**F**). **G** Dot plot showing expression of enteric mesothelial fibroblast marker genes identified in Zeisel et al., 2018 in clusters of the adult meta-atlas.

**Supplementary Figure 3:**
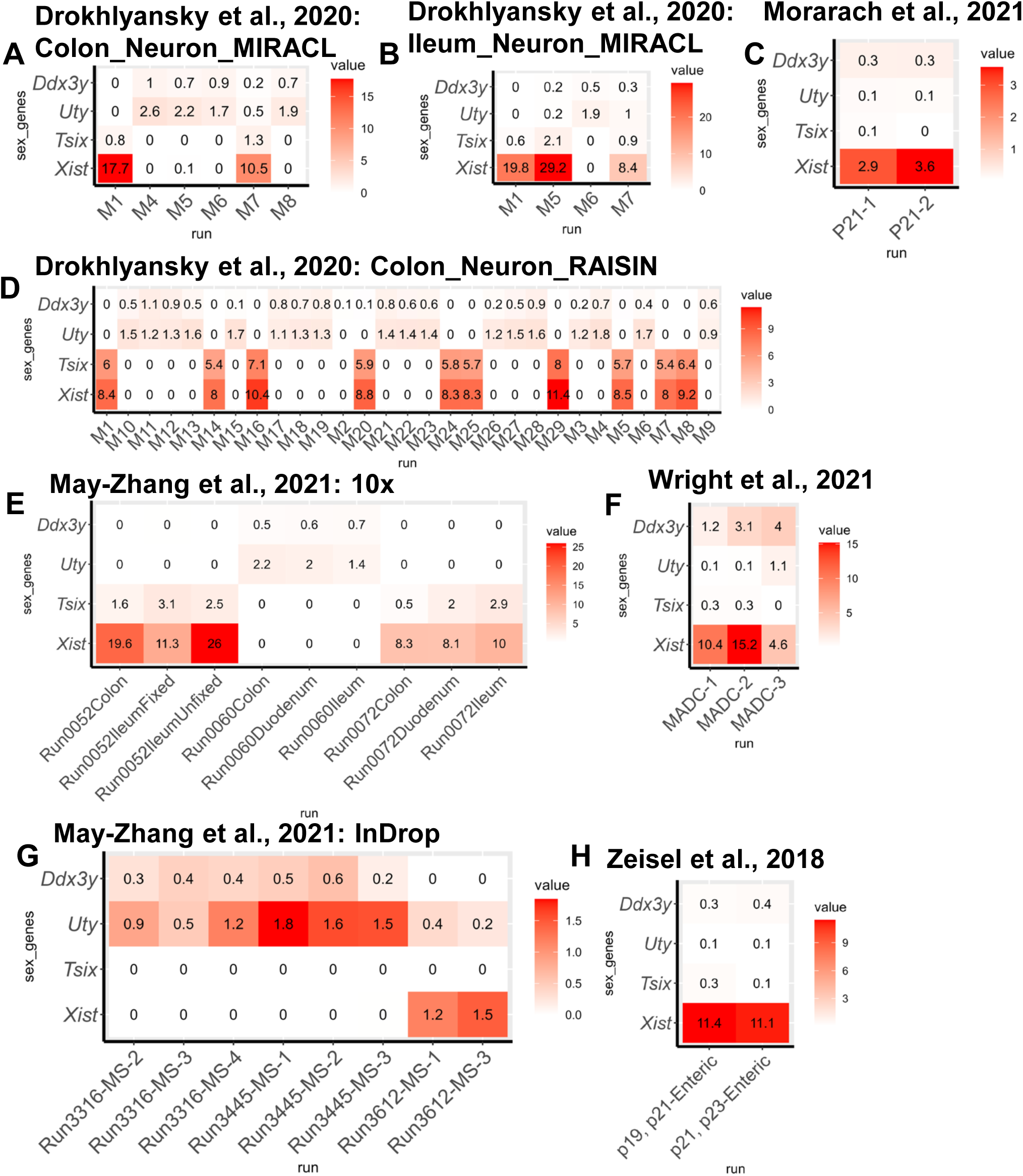
Sex-biased gene expression of *Ddx3y*, *Uty*, *Tsix*, and *Xist* in each sc/snRNA-seq component datasets. **A** Heatmap of expression of sex-biased genes in MIRACL-seq colon neuronal cells from Drokhlyansky et al., 2020 split by mouse. **B** Heatmap of expression of sex-biased genes in MIRACL-seq Ileum neuronal cells from Drokhlyansky et al., 2020 split by mouse. **C** Heatmap of expression of sex-biased genes in Morarach et al., 2021 split by scRNA-seq run. **D** Heatmap of expression of sex-biased genes in RAISIN-seq colon neuronal cells from Drokhlyansky et al., 2020 split by mouse. **E** Heatmap of expression of sex-biased genes in May-Zhang et al., 2021 10X data split by snRNA-seq run. **F** Heatmap of expression of sex-biased genes in Wright et al., 2021 split by snRNA-seq run. **G** Heatmap of expression of sex-biased genes in May-Zhang et al., 2021 InDrop data split by snRNA-seq run. **H** Heatmap of expression of sex-biased genes in Zeisel et al., 2018 split by snRNA-seq run.

